# Endolysosomal dysfunction in radial glia progenitor cells leads to defective cerebral angiogenesis and compromised Blood-Brain Barrier integrity

**DOI:** 10.1101/2023.09.24.559123

**Authors:** Ivan Bassi, Moshe Grunspan, Gideon Hen, Kishore A. Ravichandran, Noga Moshe, Stav R. Safriel, Amitai Chen, Carmen Ruiz de Almodovar, Karina Yaniv

## Abstract

The neurovascular unit (NVU) is a complex structure comprising neurons, glia, and pericytes that interact with specialized endothelial cells to maintain cerebral homeostasis and blood-brain barrier (BBB) integrity. Alterations to NVU formation and function can lead to serious forms of cerebrovascular disease, including cerebral small vessel diseases (CSVDs), a range of pathological changes of cerebral capillaries within the white matter contributing to BBB dysfunction and demyelination.

Despite the growing recognition of the pivotal roles played by neuro-vascular and glia-vascular interfaces in NVU formation and functioning, CSVD research has mainly focused on characterizing pericyte and EC dysfunction, leaving our understanding of the contribution of non-vascular cells of the brain parenchyma limited.

Here, we use a novel zebrafish mutant to delve into the intricate interplay among NVU components and demonstrate how the compromised specification of a progenitor cell population sets off a cascade of events, ultimately leading to severe cerebrovascular abnormalities. The mutation affects Scavenger Receptor B2 (scarb2)/Lysosomal Membrane Protein 2 (limp2), a highly conserved protein residing in the membrane of late endosomes and lysosomes. We find Scarb2 to be predominantly expressed in Radial Glia Cells (RGCs), a multipotent cell giving rise to neurons and glia in both zebrafish and mammals. Through live imaging and genetic manipulations, we identify impaired Notch3 signaling in RGCs and their glial progeny as the primary consequence of Scarb2a depletion and show that this disruption causes excessive neurogenesis at the expense of glial cell differentiation. We further pinpoint compromised acidification of the endolysosomal compartment in mutant cells as the underlying cause of disrupted Notch3 processing, linking for the first time Notch3 defects in non-vascular cells of the brain parenchyma to CSVD phenotypes.

Given the evolutionary conservation of SCARB2 expression and the remarkable recapitulation of CSVD phenotypes, *scarb2* mutants provide a promising framework for investigating the mechanisms governing Notch3 processing in non-vascular cells and their involvement in the onset of CSVD.

## Introduction

In recent years, the concept of the neurovascular unit (NVU) has gained prominence in the fields of neuroscience and vascular biology. The NVU refers to a complex multicellular structure within the brain comprising neurons, astrocytes, oligodendrocytes (OL), pericytes, and specialized endothelial cells (ECs), which work in concert to help maintain cerebral homeostasis by controlling blood flow and ensuring the proper functioning of the blood-brain barrier (BBB)^1,2^. ECs forming the BBB vessel wall exhibit a distinctive array of characteristics that set them apart from ECs found in other vascular beds^3,4^. Their remarkable organotypicity stems from their ability to tightly control the passage of substances between the bloodstream and the brain parenchyma due to the presence of tight junctions, which form a nearly impermeable barrier that shields the delicate neural environment from potential toxins and fluctuations in systemic composition. Recent studies have also revealed the unique gene expression repertoire of BBB-ECs^4,5^. While this EC specialization is fundamental to maintaining brain homeostasis and safeguarding neurological integrity, its underlying molecular basis has only recently begun to be elucidated.

Brain vascular abnormalities can lead to various, sometimes life-threatening forms of cerebrovascular disease^6^, including ischemic and hemorrhagic stroke, vascular malformations, vascular cognitive impairment, and dementia. Within this category, cerebral small vessel diseases (CSVDs) encompass a range of sporadic or hereditary pathological alterations of cerebral arterioles, capillaries, and venules, occurring primarily within the white matter. These changes contribute to demyelination, BBB dysfunction, glial activation, and axonal loss^7^. Despite the growing recognition of the crucial roles played by neuro-vascular and glia-vascular interfaces in the formation and functioning of the NVU and BBB, most of the research on cerebrovascular diseases and CSVD, in particular, has primarily focused on characterizing pericyte and EC dysfunction. In contrast, our understanding of how cells of the neural parenchyma contribute to CSVD and the molecular signaling pathways underlying their pathological interactions with the vasculature is poorly understood.

During embryonic development, multipotent radial glia cells (RGCs), serve as the primary source of neurons and glia (*i.e.,* astrocytes and oligodendrocytes)^8^. Oligodendrocytes (OL) - the myelinating cells of the central nervous system (CNS)-, and astrocytes, star-shaped glial cells with specialized endfeet that ensheath blood vessels, arise from RGCs and undergo proliferation, migration, and differentiation to generate the diverse population of glial cells. The timing and spatial distribution of glial cell development are tightly regulated by a combination of intrinsic genetic programs, extrinsic signals from neighboring cells, and environmental factors. Among them, the Notch signaling pathway, particularly the Notch3 receptor, has been shown to play critical roles in promoting RGC differentiation toward glial lineages^9–11^.

Here, we report the identification of a novel zebrafish mutant featuring impaired RGC differentiation and aberrant gliogenesis, which lead to defective cerebral angiogenesis, early onset of micro-hemorrhages, BBB dysfunction, and compromised myelination, all highly reminiscent of CSVD phenotypes. The mutation affects Lysosome Membrane Protein II (Limp2)/ Scavenger Receptor Class B Member 2 (SCARB2), a protein found in the membrane of late endosomes and lysosomes^12^, highly conserved in vertebrates, including humans (Extended Data Fig. 1a). Using live imaging and genetic manipulations, we identify impaired Notch3 signaling as one of the major outcomes of Scarb2a depletion in RGCs, OLs, and astrocytes and provide evidence linking this phenotype to compromised acidification of the endolysosomal compartment in mutant RGCs and glia.

Overall, our findings highlight the link between aberrant glial cell differentiation and cerebrovascular defects and demonstrate that hindering Notch intracellular processing in non-vascular cells elicits pronounced defects throughout the CNS vasculature, further impacting the formation and functionality of the BBB. Moreover, we show that Scarb2 absence results in compromised acidification of the endolysosomal compartment in RGCs and glia, thereby disrupting Notch3 processing and release of the Notch intracellular domain (NICD). Given the evolutionary conservation of SCARB2 expression in glial cells and the notable similarity of the detected phenotypes, *scarb2* mutants offer a novel framework for investigating the role of non-vascular cells of the brain parenchyma in the onset of cerebrovascular diseases and open new avenues for research into the mechanisms controlling Notch3 processing in cells of the glia lineage.

## Results

We generated *scarb2a* mutants as part of a CRISPR/CAS-based mutagenesis screen aimed at identifying genes specifically required for the formation of the CNS vasculature. The mutation induces a premature stop codon at amino acid 73, resulting in the absence of the conserved C-terminal transmembrane domain and cholesterol binding site (Extended Data Fig. 1a,b). Homozygous *larvae* die at ∼10 days post-fertilization (dpf), as opposed to heterozygous animals that reach adulthood and are viable and fertile.

At 48 hours post-fertilization (hpf), *scarb2a* mutants are readily distinguishable from their wt siblings based on their short body length and curved trunk (Fig. 1a,b). In addition, ∼25% of the mutants display intraventricular hemorrhage, primarily in the hindbrain, starting at ∼40 hpf (Fig. 1b, arrow). At 4 dpf, behavioral abnormalities previously associated with epileptic seizures, such as burst activity and whole-body trembling^13^, were also observed (data not shown).

**Fig. 1:**
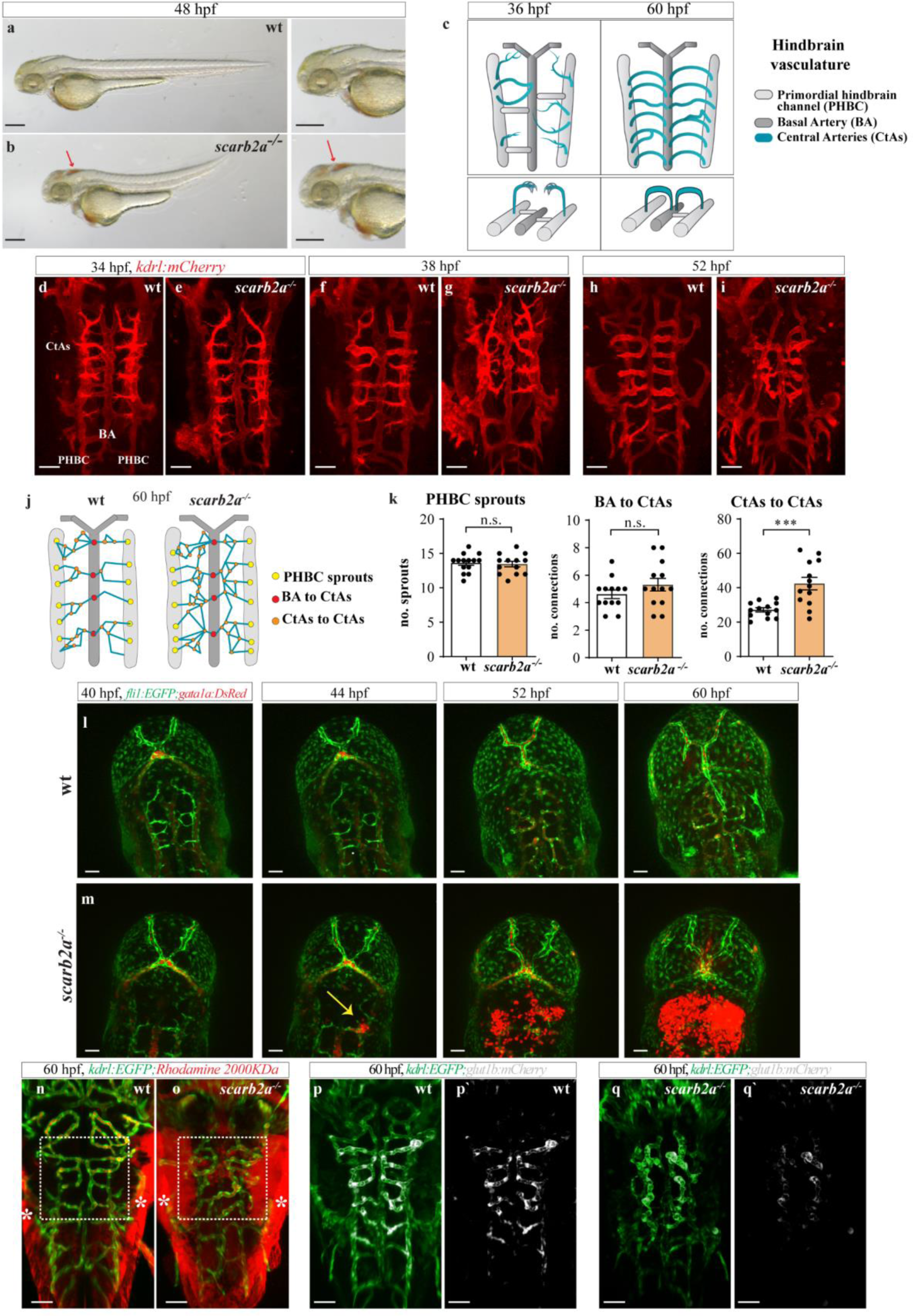
*scarb2a* mutants feature impaired cerebral angiogenesis and dysfunctional BBB. **a-b,** Brightfield images of wild-type (wt) and *scarb2a^−/−^*embryos at 48 hpf depicting morphological defects and intraventricular hemorrhage (b, arrow) in the mutants. **c**, Schematic diagram illustrating dorsal views of the developing zebrafish hindbrain vasculature and its different components, as shown in (d-i). **d-i,** Confocal images of *Tg(kdlr:mCherry)* wt and *scarb2a^−/−^* at 34 (d,e), 38 (f,g) and 52 (h,i) hpf, showing increasing defects in the morphology and patterning of mutant CtAs, starting at 38 hpf. **j,** Schematic reconstruction of wt and *scarb2a^−/−^*hindbrain vasculature at 60 hpf depicting parameters quantified in (k)**. k,** Quantification of wt and *scarb2a^−/−^* EC phenotypes at 60 hpf shows no differences in CtA sprouting from the PHBCs or CtA connection to the BA, but increased numbers of interconnections between the CtAs (n=13, two-tailed Student’s *t*-test, *P*= 0.0038). **l-m**, Selected confocal images from time-lapse series of *Tg(fli1:EGFP;gata1a:DsRed)* wt and *scarb2a^−/−^* show extravasation of red blood cells (RBCs) from the CtAs in the mutants, starting at ∼44 hpf (m, arrow). **n-o,** Intravascular injection of Rhodamine Dextran 2000 KDa at 60 hpf demonstrates compromised integrity and enhanced permeability of BBB vessels in mutant embryos (* mark unspecific dye accumulation in the skin). **p-q’**, Confocal images of *Tg(kdrl:EGFP;glut1b:mCherry)* show strong downregulation of *glut1b* expression in *scarb2a* mutants (*glut1b* in white). Scale bars, a,b=100 μm; d-i; k-l; m-n; p-q’=50 μm. CtAs, central arteries, PHBC, Primordial hindbrain channel, BA, basal artery. Error bars are mean ± s.e.m.

In order to ascertain the nature of the vascular phenotypes, we first obtained confocal images of the EC-specific reporter *Tg(kdrl:mCherry),* which revealed severe defects, particularly in the loop-shaped Central Arteries (CtAs). The CtAs arise as angiogenic sprouts from the dorsal surface of the Primordial Hindbrain Channels (PHBCs) between 32-36 hpf and gradually invade the hindbrain where they anastomose with the Basal Artery (BA)^14,15^ (Fig. 1c). At 34 hpf, no significant defects were observed in the PHBCs, BA, or CtAs of *scarb2a* mutants (Fig. 1d,e). By ∼38 hpf, however, wt CtAs establish connections with the BA, lumenize, and lose filopodia (Fig. 1f), whereas mutant CtAs are greatly disorganized, present numerous filopodia extensions, and establish abnormal interconnections (Fig. 1g). This phenotype worsens by 52 hpf, where wt hindbrains display a well-structured system of stereotypically patterned CtAs (Fig. 1h,j), while *scarb2a* mutants present a disorganized network of collapsed and hyper-branched capillaries surrounding the BA (Fig. 1i-k). Notably, these phenotypes were specific to the brain vasculature, as no defects were observed in the trunk intersegmental vessels at similar developmental stages (Extended Data Fig. 1c,d). We then proceeded to evaluate the emergence of the hemorrhagic bleeding by time-lapse imaging *Tg(fli1:EGFP;gata1a:DsRed);scarb2a^−/−^* embryos between 40-60 hpf (Fig. 1l,m and Extended Data Video 1). We found that in wt embryos, *gata1*^+^ erythrocytes remain normally confined within the hindbrain capillaries throughout the course of the experiment. In contrast, increasing hemorrhage areas are detected in the hindbrains of *scarb2a* mutants starting at ∼44 hpf (Fig. 1m, arrow), resulting in massive erythrocyte leakage from the CtAs into the ventricles. To investigate if *scarb2a* mutants also display a dysfunctional BBB, we conducted intravascular injection of 2000 kDa Rhodamine Dextran, a tracer known to be retained within the BBB of zebrafish embryos at 60 hpf^16^. As seen in Fig. 1n,o, *scarb2a* mutants presented clear extravasation of the dye from the CtAs, suggesting compromised integrity of the BBB. Moreover, the expression of Glucose transporter 1 (Glut1), a well-established marker of BBB ECs in vertebrates^17^, was markedly reduced at 60 hpf as evidenced by both *Tg(glut1b:mCherry;kdrl:EGFP)* confocal images (Fig. 1p-q’) and whole mount *in situ* hybridization (Extended Data Fig. 1e,f). Thus, *scarb2a* mutants feature defective CNS angiogenesis, accompanied by intracranial hemorrhage and a dysfunctional and leaky BBB.

Notably, the presence of Scarb2 in ECs has not been described^18^. Analysis of single-cell RNA-sequencing (scRNA-Seq) data^19^ derived from mouse developing brain and spinal cord, highlighted microglia, newly-formed OLs (NFOLs), and oligodendrocyte precursor cells (OPCs) among the top 10 cell types expressing Scarb2 (Extended Data Fig. 2a). Similarly, Scarb2 is primarily expressed by OPCs^20,21^ and microglia^22^ in humans. In zebrafish embryos, the presence of *scarb2a* mRNA was reported in the notochord and throughout the developing CNS at 24 hpf^23^. However, the specific cell type in the brain expressing *scarb2a* was not described. To accurately map *scarb2a* expression and identify the cell type/s affected by the mutation, we generated a transgenic reporter-*TgBAC(scarb2a:KalTA4)* expressing the KalTA4 driver downstream of the *scarb2a* promoter, following removal of the *scarb2a* coding sequence (Extended Data Fig. 2b). This strategy enabled visualization of *scarb2a-*expressing cells both in wt and mutant embryos. Confocal images of *TgBAC(scarb2a:KalTA4;UAS:mkate2;kdrl:EGFP)* confirmed the absence of *scarb2a* expression in *kdrl^+^* ECs at 48 (Fig. 2a-a’’) and 60 hpf (Extended Data Fig. 2c-c’’). Instead, the *scarb2a* reporter highlighted a population of cells in the ventricular zone (vz, Fig. 2a’-a’’,b) highly reminiscent of radial glial cells (RGCs), a population of multipotent cells that serve as the primary source of neurons and glia in the developing embryo^24^. RGCs display a unique elongated morphology (Fig. 2a’’’) and express specific markers, such as glial fibrillary acidic protein (GFAP)^25^ and SOX2, a transcription factor crucial for maintaining them in a progenitor state, preventing their premature neuronal differentiation^26,27^. Indeed, confocal imaging of *TgBAC(scarb2a:KalTA4;UAS:EGFP;GFAP:dTomato*) embryos revealed that at 24 hpf, a subset of *scarb2a^+^* cells is also labeled by the GFAP transgene (Fig. 2c-d’) and by Sox2 antibody^28^ (Fig. 2e-f’) in both wt and mutant embryos. Thus, based on their morphology, gene expression, and temporal distribution, we identified *scarb2a*^+^ cells as RGCs and concluded that they are unaffected by the mutation up to this developmental stage.

**Fig. 2:**
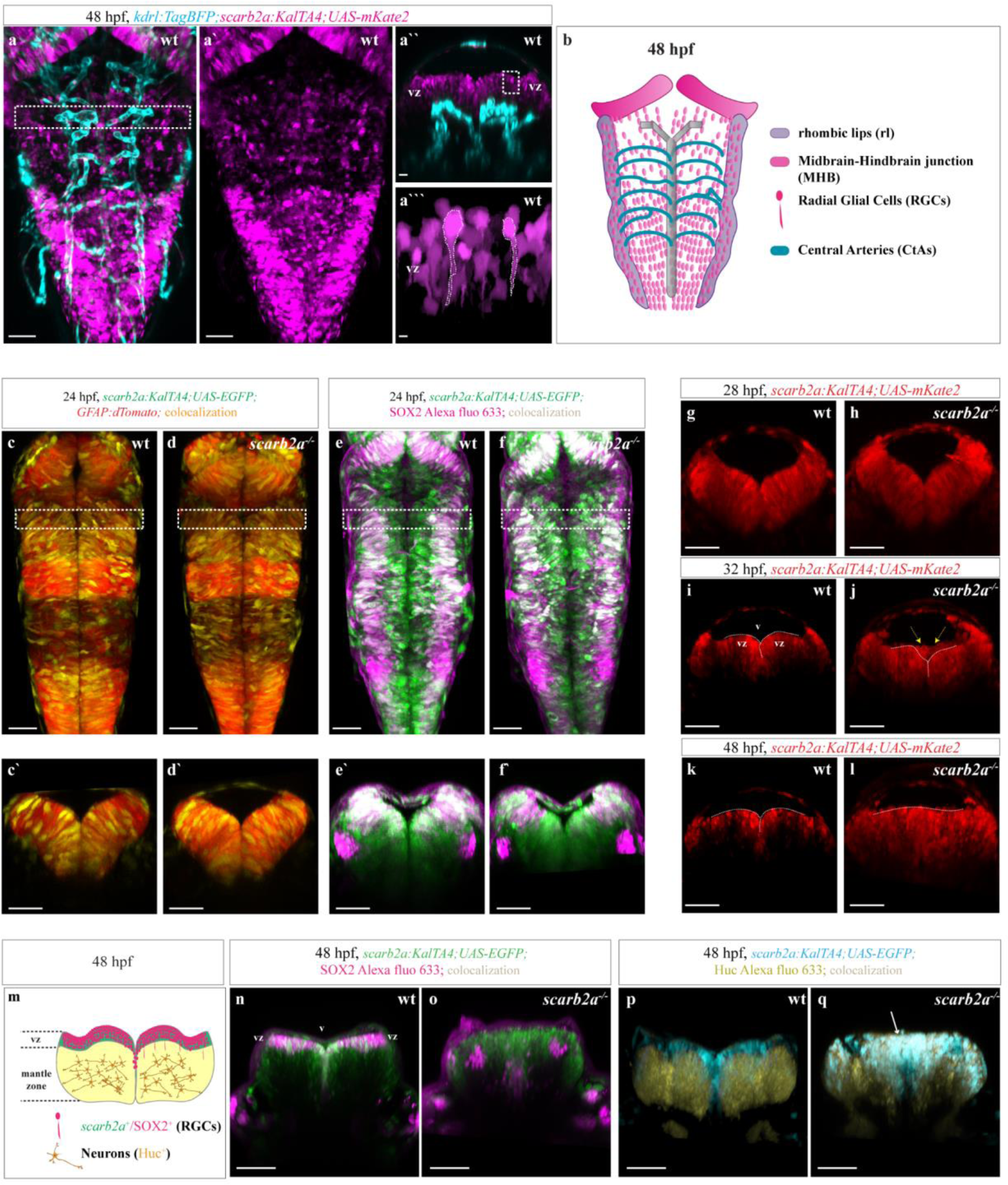
*scarb2a* is expressed in radial glial cells (RGCs), and its depletion affects brain morphogenesis. **a-a’**, Confocal images of *Tg(kdrl:TagBFP;scarb2a:KalTA4;UAS-mkate2*) show no co-expression of *scarb2a* (magenta) and the *kdrl* (blue) at 48 hpf (dashed square in (a) marks the region displayed in (a’’)). **a’’,** Transverse optical section of area shown in (a) demonstrates the presence of *scarb2a^+^*cells in the ventricular zone (vz; dashed square depicts the region enlarged in (a’’’)). **a’’’,** *scarb2a^+^* cells (dashed lines) display slender, elongated morphology typical of RGCs. **b**, Schematic illustration depicting the anatomical organization of RGCs and blood vessels in the zebrafish hindbrain at 48 hpf. **c-d’**, Dorsal views of wt and mutant hindbrains showing co-localization of *Tg(scarb2a:KalTA4;UAS-EGFP)* and *Tg(GFAP:dTomato)* signal in a subpopulation of RGCs (yellow channel denotes co-localization, dashed squares in c-d mark the level of the optical sections in c**’-**d’). **e-f’** Dorsal views of wt and mutant hindbrains showing co-localization of *Tg(scarb2a:KalTA4;UAS-EGFP)* and SOX2 immunostaining in RGCs at the vz (co-localization depicted in white, dashed squares in e-f mark the level of the optical sections in e**’-**f’). **g-l**, Selected transverse optical sections from a time-lapse series of *Tg(scarb2a:KalTA4;UAS-mkate2)* in wt (g,i,k) and mutant (h,j,l) embryos. Images show gradual restriction of *scarb2a* expression to the vz at 28 (g-h), 32 (i-j), and 48 (k-l) hpf in wt fish. In mutant embryos, *scarb2a*-labeled cells invade the ventricle space starting at 32 hpf (j-l). **m,** Schematic representation of a transverse section at the hindbrain level depicting organization of different cell types at 48 hpf. **n-o**, Immunofluorescence staining of *Tg(scarb2a:KalTA4;UAS-EGFP)* at 48 hpf showing SOX2 expression in *scarb2a^+^* RGCs in the vz of wt embryos (n, magenta), that is utterly absent in the vz of *scarb2a* mutants (o). **p-q**, HuC immunostaining on *Tg(scarb2a:KalTA4;UAS-EGFP)* embryos at 48 hpf showing HuC+ cells in the mantle zone of wt embryos (p, yellow) and scarb2+ RGCs in the vz (p, blue). In *scarb2a* mutants, *scarb2a*-labeled cells co-express HuC and are seen in the vz, mantle zone, and invading the ventricular space (q, arrow), indicating they have adopted a neuronal fate (white depicts co-localization channel). Scale bars, a-a’=50 μm; a’’=25 μm; a’’’=5 μm; c-f=50 μm; c’-f’, g-l, n-q=30 μm. vz, ventricular zone, v, ventricle space. Error bars are mean ± s.e.m.

Since the vascular phenotypes typically appear around 30 hpf, we decided to carry out time-lapse imaging of wt and mutant *TgBAC(scarb2a:KalTA4;UAS:mkate2)* embryos between 24-40 hpf to identify potential defects in the *scarb2a*^+^ cells that could lead to vascular abnormalities (Extended Data Video 2). Transverse optical sections at the hindbrain level showed no apparent defects at the initial stages of secondary neurogenesis (∼28 hpf) in *scarb2a* mutants (Fig. 2g,h). However, starting at ∼32 hpf, *scarb2a^−/−^-*labeled cells from the ventricular zone begin invading the ventricular space (v) in mutant hindbrains (Fig. 2i,j, arrows; Extended Data Fig. 2d,e). These phenotypes become more pronounced as development progresses, leading to disrupted morphology and midline and bilateral cell organization loss by 40 hpf (Fig. 2k,l; Extended Data Fig. 2f,g). At 48 hpf, we observed two well-defined domains in the hindbrain of wt embryos: the vz containing undifferentiated RGCs co-expressing *scarb2a:EGFP* and Sox2 (Fig. 2m,n) and the sub-ventricular and mantle zones, with cells labeled by the pan-neuronal marker HuC^+^ that were largely devoid of *scarb2a* expression (Fig. 2p). In contrast, there were no *scarb2a*/Sox2 double-positive cells in the vz of mutants embryos (Fig. 2o); rather most cells highlighted by the *scarb2a:EGFP* reporter were also labeled by the HuC^+^ antibody (Fig. 2q), suggesting that they have acquired a neuronal fate. Furthermore, the neuronal domain had significantly expanded, with newly differentiated neurons overtaking the vz and invading the ventricular space (Fig. 2q, arrow). Thus, our findings indicate that *scarb2a* is expressed in a subset of RGCs during early neurogenesis, and its depletion results in the loss of SOX2^+^ undifferentiated RGCs in the vz accompanied by excessive neuronal differentiation and overall abnormal hindbrain morphogenesis. Moreover, our data define a concerted temporal onset of both neuronal and vascular phenotypes.

After neurogenesis, the remaining undifferentiated RGCs generate OLs and astrocytes through asymmetric cell division^8,25,29^. The marked reduction of RGCs in *scarb2a* mutants raises the question of how this impacts gliogenesis. In the zebrafish hindbrain, Olig2-labeled OPCs emerge around 30 hpf from the pMN clusters located at the midline in rhombomeres r5 and r6^30^. To investigate whether Scarb2a depletion affects the OPC and/or OL populations, we generated *scarb2a* mutants under *olig2:EGFP* and *mbp:EGFP* transgenic backgrounds. Confocal images at 30 hpf showed no differences in the pMNs of wt and *scarb2a^−/−^* embryos (Extended Data Fig. 3a,b). By 60 hpf, when the newly generated OPCs begin to migrate to colonize the entire CNS^30^, wt *olig2*^+^ cells were found in the pMNs as well as spread radially throughout the hindbrain (Fig. 3a,c,d). In contrast, *scarb2a* mutant hindbrains exhibited significantly reduced numbers of *olig2*^+^ OPCs, both within and outside the pMNs (Fig. 3b,e,f,f’), suggesting that the mutation affects both OPC formation and migration. Mature OLs produce myelin and express myelin binding protein (MBP), a major constituent of the myelin sheath^31^. In zebrafish, myelin production begins at 3 dpf, when OPCs start maturing to become myelinating OLs^32^. Importantly, *Tg(mbp:EGFP);scarb2a^−/−^*embryos display a significant decrease in the number of myelinating OLs at 5 dpf (Fig. 3g-k), suggesting that the early oligodendrogenesis defects do not recover at later developmental stages. Interestingly, analysis of publicly available scRNA-Seq data derived from 310 cells isolated from *Tg(olig1:memEYFP),* a zebrafish reporter that specifically labels OPCs and OLs^33^, revealed the presence of *scarb2a* in the oligodendrocyte lineage (Extended Data Fig. 3c), suggesting that apart of acting on RGCs, Scarb2a could modulate additional steps during oligodendrocyte lineage progression.

**Fig. 3:**
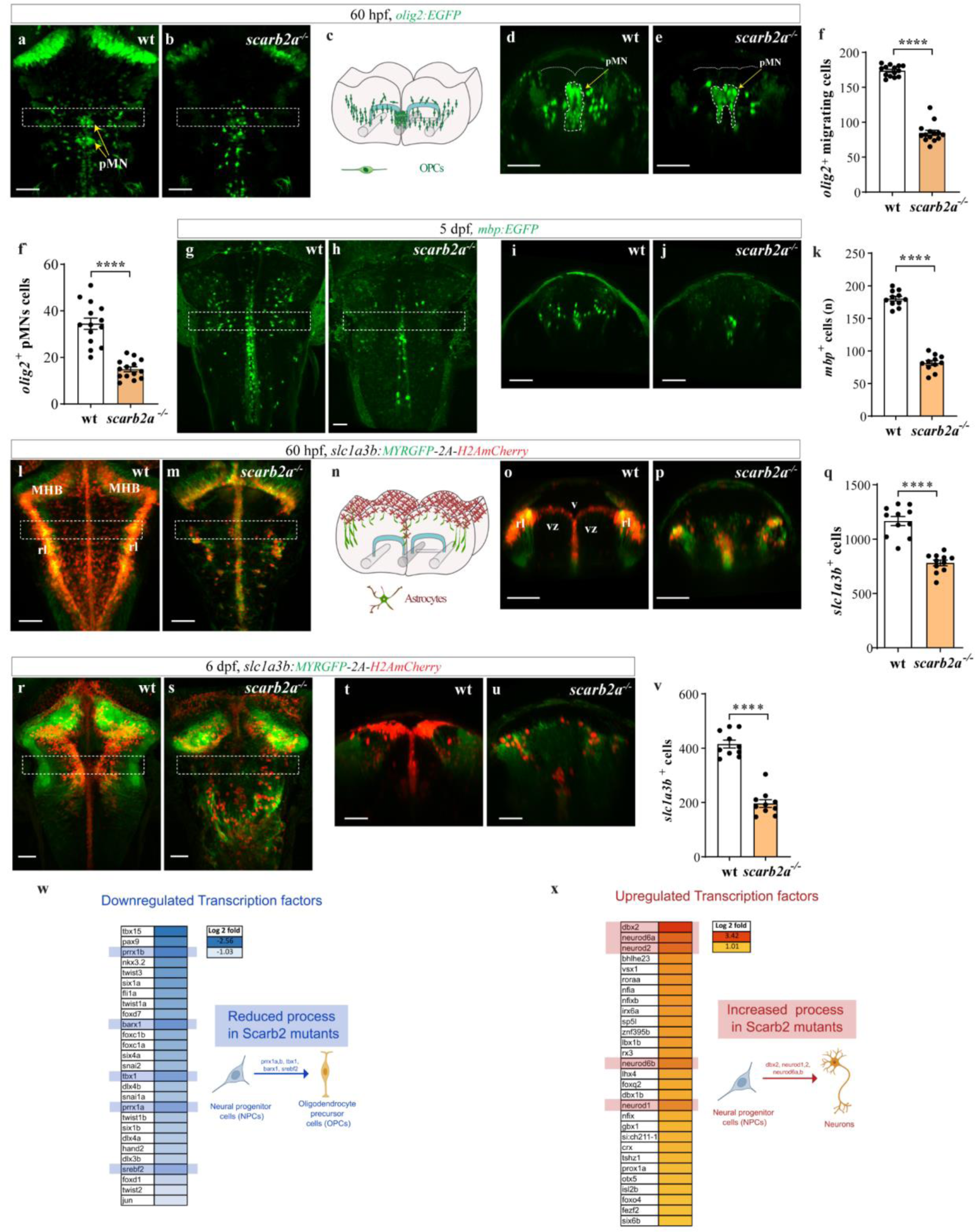
*scarb2a* depletion leads to impaired gliogenesis. **a-b,** Dorsal views of *Tg(olig2:EGFP)* wt (a) and *scarb2a^−/−^* (b) at 60 hpf showing a significant reduction of OPCs in the hindbrain of mutant embryos (dashed squares in a-b mark the level of the optical sections showed in d,e). **c,** Diagram illustrating OPC distribution in transverse section of the hindbrain at 60 hpf. **d,e** Transverse optical sections of embryos shown in (a) and (b) depicting the reduction of *olig2^+^* migrating cells and in the pMN domain (dashed lines) in *scarb2a* mutants **f,** Quantification of *olig2^+^* migrating OPCs (n=14, two-tailed Student’s *t*-test, *P*<0.0001) and **f’,** *olig2^+^* OPCs in the pMN domain (n=14, two-tailed Student’s *t*-test, *P*<0.0001). **g-k,** Dorsal views (g,h) and transverse optical sections (i,j) of *Tg(mbp:EGFP)* embryos at 5 dpf depicting reduced numbers of myelinating OLs in *scarb2a^−/−^*, quantified in (k) (n=14, two-tailed Student’s *t*-test, *P*<0.0001). **l-m,** Confocal images of *Tg(slc1a3b:MYRGFP-2A-H2AmCherry)* embryos showing fewer astrocyte nuclei in mutant hindbrains at 60 hpf (dashed squares in l-m mark the level of the optical sections shown in o,p). **n**, Graphical representation of astrocyte distribution in transverse sections of wt hindbrain at 60 hpf. **o-q** Transverse optical section of embryos shown in (l) and (m) showing absence of *slc1a3b^+^* red nuclei in the vz of *scarb2a* mutants, quantified in (q) (n =11, two-tailed Student’s t-test, P<0.0001). **r-v** Dorsal views (r,s) and transverse optical sections (t,u) of *Tg(slc1a3b:MYRGFP-2A-H2AmCherry)* embryos at 6 dpf showing reduced numbers of astrocytes, quantified in (v) (n=10, two-tailed Student’s t-test, P<0.0001). **w,x**, RNA-seq data analysis showing down (w) and up-regulated (x) TF expression in *scarb2a* mutant embryos, identified using Uniprot gene ontology. pMN= primary motor neuron, MHB= midbrain-hindbrain junction, rl= rhombic lips, vz=ventricular zone. Scale bars, a-b, g-h, l-m, r-s=50 μm; d-e, i-j, o-p, t-u =30 μm. Error bars are mean ± s.e.m.

Astrocytes also arise from RGCs^34^. To evaluate whether astrogenesis is also impaired in *scarb2a* mutants, we imaged *Tg(slc1a3b:MYRGFP-2A-H2AmCherry)* embryos that express the membrane-bound myristoyl-GFP (myrGFP) and the nuclear marker H2AmCherry under the control of *slc1a2b/Glast* promoter, a well-established astrocyte marker^35^. Confocal images at 60 hpf showed the presence of *slc1a3b^+^* cells throughout the hindbrain, particularly at the midline, midbrain-hindbrain junction (MHB), and rhombic lips (rl) (Fig. 3l). Transverse optical sections indicated that the nuclei of these cells (red) are situated in the vz and the rhombic lips, from where long GFP^+^ processes extend ventrally into the neural tube (Fig. 3n,o). Moreover, images of double transgenic *Tg(slc1a3b:MYRGFP-2A-H2AmCherry; scarb2a:KalTA4:UAS:EGFP)* embryos depicted *scarb2a*-EGFP^+^ and *slc1a3b*-mCherry^+^ colocalization in the vz nuclei, indicating that *scarb2a* is also expressed in this cell population (Extended Data Fig. 3d,d’). As opposed to wt siblings, *scarb2a^−/−^* embryos exhibited significantly fewer astrocyte nuclei at all locations (Fig. 3m,p,q), especially in the vz. These defects persisted through 6 dpf (Fig. 3r-v) when astrocytes become functional^36^.

To understand the genetic programs linked to the observed phenotypes, we performed bulk RNAseq on *scarb2a*^+^ cells isolated from wt and mutant heads at 60 hpf. Analysis of transcription factor (TF) expression revealed a significant reduction of factors involved in OPC differentiation^37,38^ (Fig. 3w), accompanied by a concomitant increase in TFs driving neuronal fate^39^ (Fig. 3x) in *scarb2a^−/−^*mutants. These results were further validated by examining *Tg(isl1a:GFP)* embryos, which revealed a marked increase in the number of differentiated motoneurons^40^ in the hindbrain of *scarb2a* mutants as compared to their wt siblings, both at 60 hpf and at 6 dpf (Extended Data Fig. 3e-j). Taken together, our data indicate that Scarb2a depletion results in precocious neuronal differentiation of RGCs and exhaustion of the progenitor pool, ultimately leading to impaired gliogenesis.

We speculated that the defects in RGCs and/or the consequent imbalanced neuronal-glia differentiation are responsible for the observed vascular phenotypes. To test this hypothesis, we conducted rescue experiments by expressing the wt *scarb2a* coding sequence in GFAP^+^ cells. Injection of a *UAS:scarb2a-P2A-RFP* or *UAS:scarb2a;cmcl2:EGFP* constructs into Tg*(GFAP:Gal4FF);scarb2a^−/−^* embryos, fully rescued the number and distribution of OPCs (Fig. 4a-d; Extended Data Fig. 3k-m) and astrocytes (Fig. 4e-h; Extended Data Fig. 3n-p) at 60 hpf. In line with our hypothesis, restoring proper gliogenesis resulted in significant recovery of CtA morphology, reduced numbers of ectopic interconnections (Fig. 4i-l), and rescued *glut1b* expression to wt levels (Fig. 4m-o’). Similar results were obtained upon expression of wt *scarb2a* in Tg*(scarb2a:KalTA4);scarb2a^−/−^* embryos (Extended Data Fig. 4a-o’), confirming that *scarb2a* depletion in RGCs prevents proper glia differentiation and maturation, leading to subsequent defects in the establishment of the hindbrain vasculature and the BBB.

**Fig. 4:**
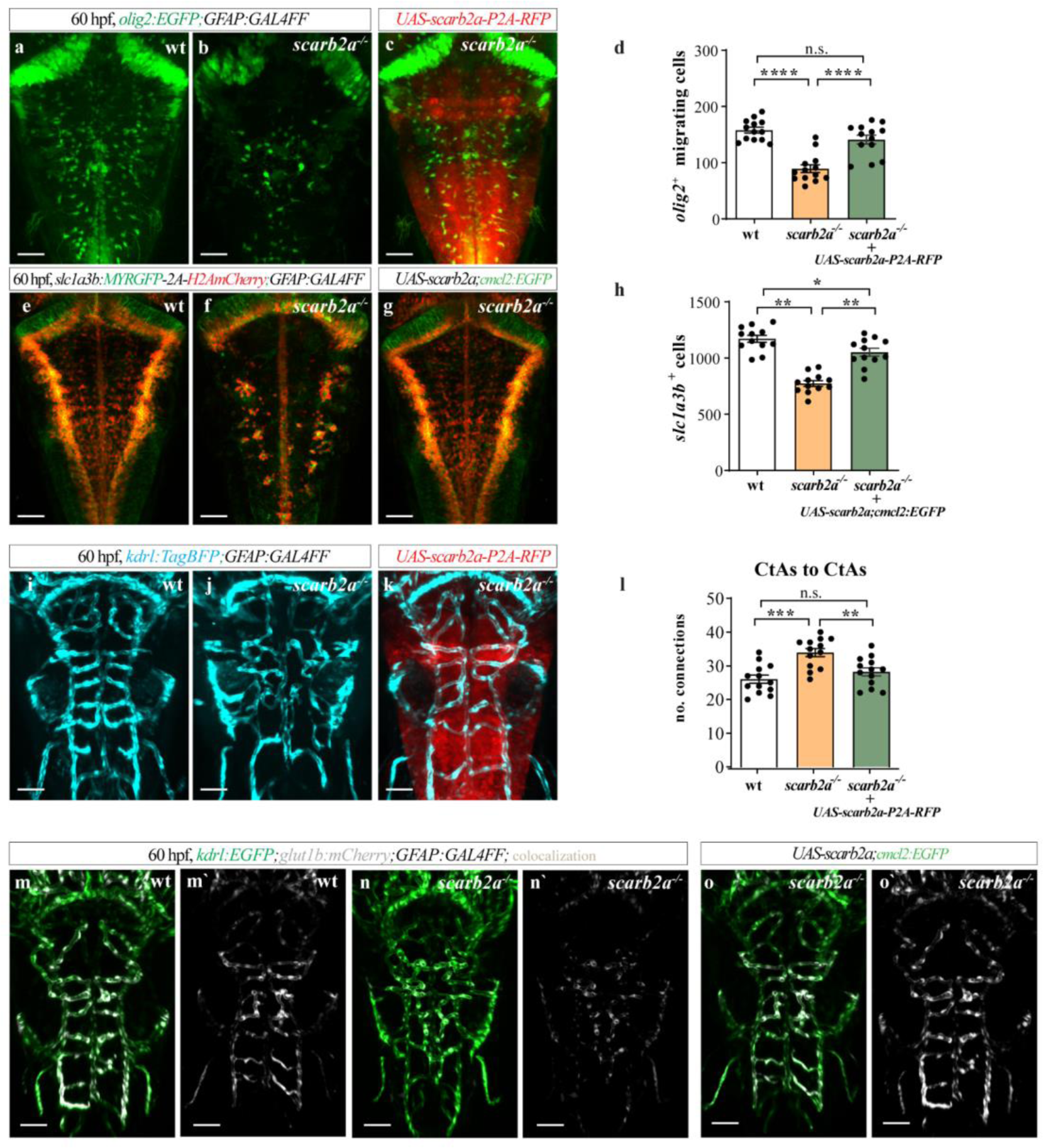
Restoring *scarb2a* expression in RGCs fully rescues the mutant phenotypes. **a-c**, Dorsal views of *Tg(olig2:EGFP;GFAP:Gal4FF)* hindbrain in wt (a), *scarb2a^−/−^* (b), and *scarb2a^−/−^* mutant following injection of *UAS-scarb2a-P2A-RFP* construct (c). **d**, Quantification of migrating *olig2^+^*cells (n=13; One-way ANOVA, multiple comparisons with Tukey posthoc test). **e-g**, Dorsal views of *Tg(slc1a3b:MYRGFP-2A-H2AmCherry; GFAP:Gal4FF)* hindbrains in wt (e), *scarb2a^−/−^* (f) and *scarb2a^−/−^* mutant following injection of *UAS-scarb2a,cmcl2:EGFP* construct (g). **h**, Quantification of *slc1a3b*^+^ red nuclei (n=12; One-way ANOVA, multiple comparisons with Tukey posthoc test). **i-k**, Dorsal views of the hindbrain vasculature in *Tg(kdrl:TagBFP;GFAP:Gal4FF)* embryos wt (i), *scarb2a^−/−^* (j) and *scarb2a^−/−^* mutant following injection of *UAS-scarb2a-P2A-RFP* construct (k). **l**, Quantification of CtA interconnections (n= 13; One-way ANOVA, multiple comparisons with Tukey posthoc test). **m-o’**, Dorsal views of *Tg(kdrl:EGFP;glut1b:mCherry;GFAP:Gal4FF)* depicting the full restoration of *glut1b* expression (white) following injection of *UAS-scarb2a,cmcl2:EGFP* construct in mutant embryos (o,o’). Scale bars, a-c; e-g; i-k; m-o’= 60 μm. Error bars are mean ± s.e.m.

The pronounced neurogenic phenotype observed in *scarb2a* mutants resembled the one resulting from Notch deficiency^41–43^, raising the possibility that *scarb2a* depletion hinders Notch signaling, resulting in precocious neuronal differentiation of RGC progenitors followed by impaired glia differentiation and maturation. To test this hypothesis, we time-lapse imaged Notch signaling during different phases of neurogenesis using the *Tg(EPV.Tp1-Mmu.Hbb:EGFP)^ia12^* (a.k.a *12NRE:EGFP* ^44^) Notch reporter under wt and *scarb2a* mutant backgrounds (Extended Data Video 3). Analysis of transverse optical sections revealed comparable levels of Notch activity in the rl of both wt and mutant embryos during the initial phase of secondary neurogenesis (28 hpf) (Fig. 5a-b’; Extended Data Fig. 5a,b). However, while at 32 hpf, vz progenitors display evident Notch activity in wt embryos (Fig. 5c,c’; Extended Data Fig. 5c), the EGFP signal was barely detected in these cells in mutant siblings (Fig. 5d,d’; Extended Data Fig. 5d). By 40 hpf, only residual levels of Notch-derived EGFP signal were observed in the rl and midline of *scarb2a* mutants (Fig. 5e-f’; Extended Data Fig. 5e,f). These results were further validated by qRT-PCR analyses carried out on *Tg(scarb2a:KalTA4;UAS:mKate2)* cells isolated from the heads of wt and *scarb2a^−/−^* embryos. The expression of the Notch downstream targets *her4.2* and *her6* was significantly reduced (Fig. 5g), whereas no differences were observed in the levels of *notch1a* and *notch3* (Fig. 5h), indicating that Scarb2a depletion impairs Notch signaling without affecting transcription of the receptors.

**Fig. 5.**
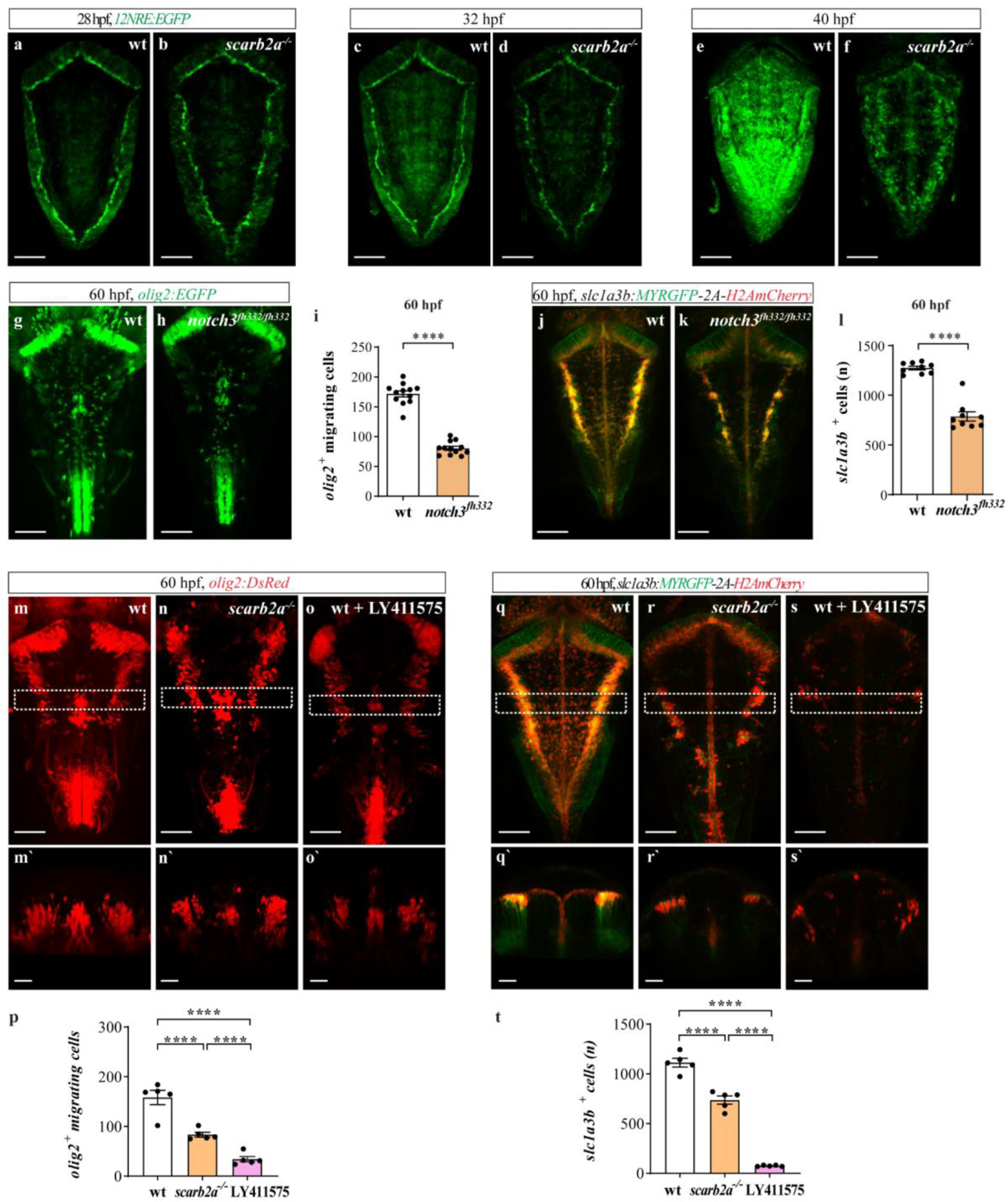
Impaired endolysosomal acidification results in Notch signaling inhibition in *scarb2a* mutants. **a-f**’ Optical transverse sections from a time-lapse series of *Tg(12NRE:EGFP;scarb2a:KalTA4;UAS-mkate2)* wt and mutant embryos at 28 (a-b’), 32 (c-d’) and 40 hpf (e-f’) showing reduced Notch signaling in *scarb2a* mutant vz starting from 32 hpf (d,d’; f,f’). **g-h** qRT-PCR analyses showing mRNA levels of *her4.2*, *her6* (g) and *notch1a, notch3* (h) in *mKate2^+^* cells sorted from *Tg(scarb2a:KalTA4;UAS-mkate2)* heads (N=5, n=20, two-tailed Student’s *t*-test, *P_her4.2_*=0.0017; *P_her6_*=0.0012). **i-j’** Transverse optical sections of *Tg(12NRE:EGFP;scarb2a:KalTA4;UAS-mkate2)* wt embryos untreated (i,i’) or treated with v-ATPase inhibitor Bafilomycin A1 (BafA1, j,j’) showing intraventricular RGC invasion (j, arrow) and reduced Notch signaling in treated embryos. **k-m.** Lysotracker Deep Red staining of *Tg(hsp70l:lamp1-RFP)* wt (k-k’’) and *scarb2a* mutant (l-l’’) embryos showing reduction of *Lamp1^+^* acidic punctae in mutant hindbrains; quantified in (o). **m-n’.** Transverse optical sections of *Tg(hsp70l:lamp1-RFP)* wt embryos treated with BafA1 (m,m’’,o) or with the γ-secretase inhibitor (LY-411575) (n,n’). **p.** Quantification of *Lamp1^+^* vesicle size in wt, *scarb2a* mutants, BafA1, and LY-411575 wt-treated embryos (for each group n=5, One-way ANOVA, multiple comparisons with Tukey posthoc test). **q-r’** Lysotracker Deep Red staining of *Tg(h2afx:EGFP-RAB7)* wt (q-q’’) and *scarb2a* mutant (r-r’’) embryos showing reduction of double positive *Rab7*/Lysotracker punctae in mutant hindbrains; quantified in (u). **s-t’**. Transverse optical sections of *Tg(h2afx:EGFP-RAB7)* wt embryos treated with BafA1 (s-s’’,u) or LY-411575 (t,t’). **v** Quantification of Rab7 vesicle size in wt, *scarb2a* mutants, BafA1, and LY-411575 wt-treated embryos (from each group n=5, One-way ANOVA, multiple comparisons with Tukey posthoc test). Scale bars, a-f’, i-j’,k,l,m,n,q,r,s,t=30 μm; k’,l’,m’,n’,q’,r’, s’,t’= 5 μm. g,h,o,u= Error bars are mean ± s.e.m.; p,v= bars represent median

Notch1a signaling is active during early neurogenesis (18-24 hpf), whereas Notch3 is the predominant isoform acting at the stages when the *scarb2a* phenotypes become evident (30-55 hpf)^28,45,46^. Accordingly, *notch3^fh332/fh332^* mutants display defective oligodendrogenesis^45^ (Extended Data Fig. 5g-i) as well as premature neuronal differentiation^28^, highly reminiscent of the *scarb2a^−/−^* phenotypes. Similarly, the *Tg(slc1a3b:MYRGFP-2A-H2AmCherry*) astrocyte population was also significantly reduced in *notch3^fh332/fh332^* embryos (Extended Data Fig. 5j-l), further reinforcing the link between Scarb2 absence, Notch3 signaling deficiency and impaired gliogenesis.

In both mice and humans, Scarb2 is found in the membrane of late endosomes and lysosomes, where it was shown to fulfill several functions^12^. Therefore, we speculated that alterations to Scarb2 might disrupt the trafficking and/or processing of Notch3 within the endolysosomal compartment. Previous research in Drosophila, zebrafish, and human cells has established a connection between endolysosomal activity and Notch signaling^47,48^. In particular, it has been shown that the acidity of the endolysosomal compartment, via the V-ATPase proton pump activity, can significantly affect the efficiency of the γ-secretase-dependent cleavage of the NICD in the receiving cell, thereby altering proper Notch signaling^49–52^.

As a first step toward testing our hypothesis, we exposed 32 hpf *Tg(12NRE:EGFP, scarb2a:KalTA4;UAS:mKate2)* wt embryos to Bafylomycin A1 (BafA1), a well-established V-ATPase inhibitor. As seen in Fig. 5i-j’, the treatment led to a drastic reduction of Notch signaling^47^ and increased numbers of *scarb2a*^+^ cells invading the ventricular space, strongly resembling the phenotypes observed in *scarb2a* mutants. Next, we asked whether the acidity of the endolysosomal compartment is indeed affected in *scarb2a* mutants. To answer this question, we took advantage of *Tg(h2afx:EGFP-rab7)^mw7^* embryos, ubiquitously expressing an EGFP-fused form of the late endosome marker Rab7^53^ and *Tg(hsp70l:lamp1-RFP)^pd1064^* fish, expressing an RFP-tagged form of the Lysosomal Associated Membrane Protein 1 (Lamp1) under the heat-shock promoter^54^. We combined these transgenic reporters with *scarb2a* mutant embryos and stained them with far-red-coupled Lysotracker, which stains acidic organelles. High-magnification images of the vz demonstrated that *scarb2a^−/−^* embryos exhibited fewer numbers of both Lamp1/Lysotracker (Fig. 5k-l’’,o) and Rab7/Lysotracker (Fig. 5q-r’’,u) double-positive punctae, as compared to wt siblings, confirming the imapired acidity of these intracellular compartments. Similar, albeit more pronounced results, were obtained upon exposure of *Tg(hsp70l:lamp1-RFP)* and *Tg(h2afx:EGFP-rab7)* wt embryos to BafA1-in this case, no punctate Lysotracker staining was detected (Fig 5m-m’’,o,s-s’’,u) further validating the notion that *scarb2a* mutants display impaired acidification of Rab7 and Lamp1 organelles. Interestingly, we also noticed that both Rab7 and Lamp1 vesicles were significantly enlarged in the mutants (Fig. 5p,v), possibly due to the accumulation of undegraded materials as has been shown in the context of several Lysosomal Storage Diseases (LSDs)^55^, or as a result of defective fusion.

Two lines of evidence confirmed that impaired S3 cleavage of Notch3 and generation of the NICD are among the main drivers of *scarb2a* mutant phenotypes. First, inhibition of γ-secretase via LY-411575 resulted in a marked reduction in the numbers of OPCs (Extended Data Fig. 5m-p) and astrocytes (Extended Data Fig. 5q-t) as well as in enlarged Lamp1 and Rab7 organelles fully phenocopying the *scarb2a*^−/−^ phenotypes (Fig. 5p,v). Second, we designed rescue experiments based on overexpression of the Notch3 intracellular (N3ICD) or extracellular (N3ECD) domains. To this end, we utilized stable transgenic reporters-*Tg(hsp70I:N3ICD-EGFP)* and *Tg(hsp70I:N3ECD-EGFP)*^56^ expressing constitutively active forms of N3ICD and N3ECD under the control of the heat shock promoter *hsp70I*, and induced their expression at ∼ 28-30 hpf, mimicking the endogenous expression of Notch3^28^. As seen in Fig. 6a-c’,e, N3ECD-overexpression in *olig2:GFP;scarb2a^−/−^* embryos did not revert defective oligodendrogenesis, as opposed to a significant recovery of the number of *olig2*^+^ migrating cells detected upon N3ICD induction (Fig. 6d-e). Similar results were obtained regarding the astrocyte population-namely, overexpression of N3ICD, but not of N3ECD, led to a significant increase in the number of *slc1a3b:MYRGFP-2A-H2AmCherry^+^* cells in the hindbrains of *scarb2a^−/−^* embryos (Fig. 6f-j). Finally, we could confirm that these phenotypes are primarily attributed to defective Notch3 signaling, as overexpression of the N1aICD fragment, using *Tg(hsp70l:myc-notch1a-intra;cryaa:Cerulean)*^57^ was not sufficient to restore neither astrocyte (Extended Data Fig. 6a-d) or OPC (Extended Data Fig. 6e-h) populations in *scarb2a* mutants.

**Fig. 6.**
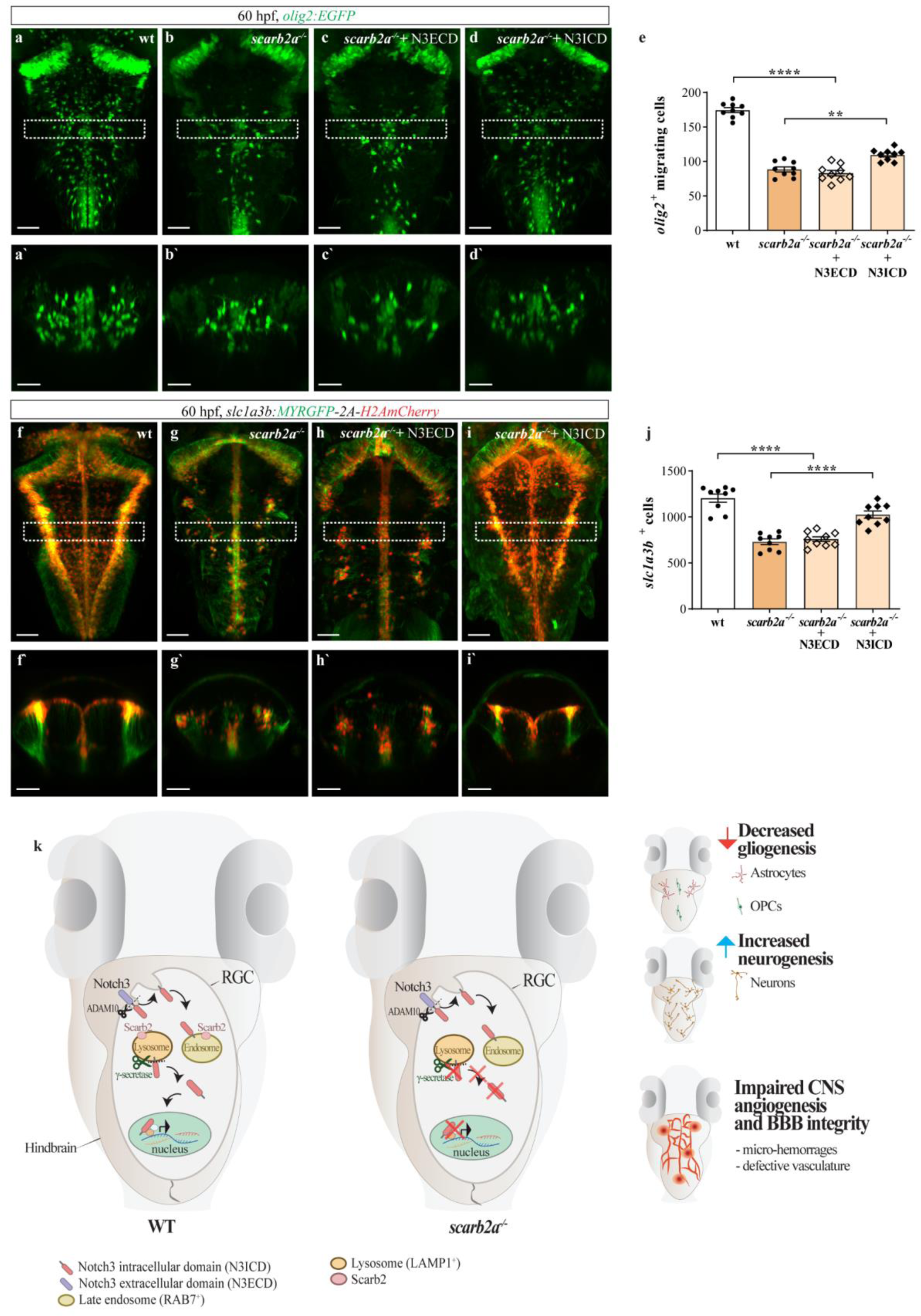
Overexpression of Notch3 intracellular (N3ICD) but not extracellular (N3ECD) domain rescues gliogenesis in *scarb2a* mutants. **a-d’** Dorsal views and transverse optical sections of *Tg(olig2:EGFP)* hindbrain in wt (a,a’), *scarb2a^−/−^* (b,b’) and *scarb2a^−/−^* following heat-shock induced overexpression of Notch3 extracellular (N3ECD) (c-c’) and Notch3 intracellular (N3ICD) domains (d-d’) (dashed square in a-d, marks the region shown in a’-d’). **e**, Quantification of migrating OPCs in a-d (n=9, One-way ANOVA, multiple comparisons with Tukey post-hoc test). **f-I’** Dorsal views and transverse optical sections of *Tg(slc1a3b:MYRGFP-2A-H2AmCherry)* hindbrains in wt (f), *scarb2a^−/−^* (g) and *scarb2a^−/−^* mutant following N3ECD (h-h’) or N3ICD (i-i’), heat-shock induction (dashed square in f-i, marks the region shown in f’-i’) **j**, Quantification of *slc1a3b^+^* astrocytes in f-i (n=9, One-way ANOVA, multiple comparisons with Tukey posthoc test). **k,** Graphical model depicting the series of events affecting NVU formation and function in *scarb2a* mutant embryos. Defective endolysosomal acidification hinders Notch3 processing in RGCs, leading to impaired gliogenesis, defective cerebral angiogenesis, micro-hemorrhages, and compromised BBB. Scale bars, a’,b’,c’,d’,f’,g’,h’,I’= 30 μm; a,b,c,d,f,g,h,i= 60 μm. Error bars are mean ± s.e.m.

Taken together, our results demonstrate that dysfunction of the endolysosomal compartment caused by Scarb2 depletion hinders Notch3 signaling in RGCs, thereby affecting the maintenance of the undifferentiated RGC progenitor pool and modulating the balance between neurogenesis and gliogenesis. We further demonstrate that alterations to this balance result in severe vascular defects, hemorrhages, and a leaky BBB (Fig. 6k). In a broader sense, our data illustrate the exquisite interactions between different cell types contributing to the establishment and functionality of the NVU and shed light on a previously overlooked relationship between Notch3 activity in non-vascular cells of the brain parenchyma (i.e., RGCs and glia), and cerebrovascular health.

## Discussion

The study of zebrafish *scarb2a* mutants has revealed new insights into the mechanisms controlling neuronal vs. glial differentiation of RGC progenitors that, when disrupted, lead to phenotypes associated with cerebrovascular diseases and demyelinating neuropathies. By generating a *scarb2a* transgenic reporter, we revealed the specific expression of *scarb2a* initially in RGCs, and subsequently in glial cells, including OPCs, OLs, and astrocytes. The absence of Scarb2 results in impaired acidification of the endolysosomal compartment in RGCs, which in turn affects the cleavage of Notch3 and the release of N3ICD. Consequently, *scarb2a* mutants display excessive neuronal formation at the expense of glial cells. This imbalance results in severe cerebrovascular abnormalities, including abnormal interconnections of the CtAs, micro-hemorrhages, and a compromised BBB.

Notably, a complete rescue of the glial and vascular abnormalities associated with Scarb2a deficiency was achieved through the expression of the wt form of *scarb2a,* exclusively in RGCs, underscoring the specificity of the observed phenotypes and the significant connection between proper RGC differentiation, gliogenesis, and CNS angiogenesis.

Previous studies have demonstrated that RGCs and their derived lineages of neurons, oligodendrocytes, and astrocytes control different aspects of embryonic and postnatal CNS angiogenesis. During the development of the embryonic mouse cortex, for instance, Wnt ligands secreted by RGCs control Glut-1 and Gpr124 expression, and their depletion results in defective CNS angiogenesis and compromised BBB integrity^58–60^. Additionally, recent research has demonstrated that RGCs secrete TGF-β1, which promotes both EC migration and tube formation *in vitro*^61^. Finally, vascular endothelial growth factor A (VEGFA), secreted by neural and/or glial cells and their progenitors, is known to control CNS angiogenesis in a dose-dependent manner^62,63^. Our results, showing clear glut1 downregulation and defective sprouting and migration of CNS ECs in mutant embryos, fully correspond with these previous reports, supporting the evolutionary conservation of these processes. In the future, it will be interesting to investigate the specific signals emanating from RGCs and their derivatives that control the development of the CtAs and sealing of the BBB in zebrafish embryos, and whether those are conserved as well.

At the molecular level, we show significant downregulation of Notch signaling in *scarb2a* mutant hindbrains starting from 30 hpf and demonstrate that this reduction is specifically attributed to impaired Notch3 activity. Through a series of genetic manipulations, we demonstrate that administration of the Notch3-intracellular, but not of the extracellular domain, leads to a recovery of OPC and astrocyte populations. These results confirm earlier findings supporting the crucial role of Notch3 in regulating RGC fate determination^11,28,64^, and highlight Scarb2a and the proper function of the endolysosomal compartment as key players in the process.

Analysis of N3ECD and N3ICD overexpression experiments showed that only the intracellular fragment is able to rescue the *scarb2a* mutant phenotype, suggesting that lack of Scarb2a might potentially affect the cleavage operated by the γ-secretase complex. The activation of the Notch signaling pathway involves two sequential proteolytic cleavages of the Notch receptor that generate first a Notch extra-cellular domain (NECD) and, subsequently, a Notch intracellular domain (NICD) fragment. The NECD is internalized within the signaling cell, whereas the product of the second (S2) cleavage, consisting of the transmembrane and intracellular domains (also referred to as Notch Extracellular Truncation-NEXT), undergoes an additional cleavage by γ-secretase at the S3 site, releasing the NICD, which can then translocate into the nucleus and activate transcription of downstream targets^65^. Experiments carried out in Drosophila^66,67^ as well as in mammalian cells^51,68^ have demonstrated that the acidity of the endolysosomal compartment can significantly affect the efficiency of the S3 cleavage (following NEXT endocytosis). Notably, inhibition of vacuolar H^+^-ATPase (V-ATPase) (a protein involved in the acidification of endosomes and lysosomes) by bafilomycin-A1 (BafA1)^66,67^ was shown to hinder the processing and activation of Notch receptors^70^, further supporting the notion that Notch S3 cleavage by γ-Sec requires an acidified intracellular compartment and that this step is crucial for Notch activation. Given that SCARB2/LIMP2 is predominantly located in the membrane of late endosomes and lysosomes, we speculated that impaired production of N3ICD observed in *scarb2a* mutants could be linked to alterations in these organelles. In this context, we show that Scarb2a depletion results in reduced acidity and increased size of both Lamp1 and Rab7 compartments, features associated with several LSDs. These data, along with the strong similarity observed between *scarb2a* mutant phenotypes and wt embryos treated with the V-ATPase inhibitor BafA1 or with the γ-Secretase inhibitor LY-411575, reinforce the notion that Scarb2-dependent malfunction of the endolysosomal machinery stands at the basis of the Notch3 signaling defects observed in *scarb2a^−/−^* mutants. Accordingly, Scarb2a depletion could lead to changes in the endolysosomal machinery and γ-secretase activity, resulting in altered N3ICD cleavage^1,71^.

In the brain, astrocytes, oligodendrocytes, and neurons exist within a microenvironment comprising various cell types, including ECs, pericytes, perivascular fibroblasts, and vascular smooth muscle cells (vSMCs) forming the neurovascular unit (NVU)^72^. Together, these cells play crucial roles in vascular-dependent brain pathologies such as CSVD and brain vascular malformations, which are characterized by the formation of leaky, tortuous, and dysfunctional vessels. Notably, CADASIL, the most common inherited form of CSVD, is caused by mutations in the *NOTCH3* gene^72^, which have been shown to primarily affect vascular smooth muscle cells and pericytes. One of the outstanding open questions is how the vascular phenotype is linked to the white matter changes observed in CADASIL (and in SVDs in general). While the current prevalent idea is that the initial vascular defects lead to white matter damage, the possibility exists that impaired Notch3 processing and/or signaling within glia, cell autonomously account for the observed changes. This hypothesis could also explain the fact that despite the monogenic nature of the disease, CADASIL patients display a large phenotypic variability and an unclear genotype-phenotype correlation.

Taken together, our findings bring about a fresh perspective on cerebrovascular diseases, suggesting that defects in cellular populations within the vascular microenvironment, beyond the commonly studied vascular cells, also play a crucial role. Specifically, our findings highlight the importance of proper endolysosomal function in less-explored cell types like RGCs and cells of the glial lineage. Interestingly, we observed that when lysosomal function is compromised in these cell types, it leads to phenotypes resembling CSVD, shedding light on a previously overlooked relationship between these cells and cerebrovascular health. Our study opens up new avenues for therapeutic approaches targeting lysosomal dysfunction in these unconventional cell players to mitigate CSVD-associated manifestations and promote better cerebrovascular function.

## Acknowledgments

The authors would like to thank H. Hasid, L. Shen, and Y. Yogev for technical assistance, K. Monks (Vollum Institute, US), C. Burns (Harvard Institute, US), B. Appel (University of Colorado, US), M. Bagnat (Duke Cancer Institute, US) and B. A. Link (Wisconsin Medical School, US) for providing reagents. S. Federici (Weizmann Institute, Israel) for assistance with statistical analysis and illustrations, G. Almog, R. Hofi, A. Glozman, and R. Brihon (Weizmann Institute, Israel) for superb animal care. The authors are grateful to all the members of the Yaniv lab for discussion, technical assistance, critical reading of the manuscript, and continuous support. This work was supported in part by Minerva Foundation (714447) to KY, ERA-Net-NEURON (Tackle-CSVD) to KY, ERC StG (LIPintoECtion 335605) to KY, and research grants from The M. Judith Ruth Institute for Preclinical Brain Research (P137071), the Weizmann – Nella and Leon Benoziyo Center for Neurological Diseases and the Weizmann SABRA - Yeda-Sela - WRC Program, the Estate of Emile Mimran, and The Maurice and Vivienne Wohl Biology Endowment and the Brenden-Mann Women’s Innovation Impact Fund. K.Y. is the incumbent of the Enid Barden and Aaron J. Jade Professorial Chair. C.R.A. acknowledges support from ERA-Net-NEURON (Tackle-CSVD) and ERC CoG (OLI.VAS 864875). I.B. was supported by a postdoctoral fellowship by the Sergio Lombroso Program and a Senior Postdoc Fellowship by the Weizmann Institute. N.M. was supported by research grants from the Estate of Olga Klein Astrachan and the Estate of Mady Dukler.

## Author contributions

I.B. designed and conducted experiments, analyzed data, and co-wrote the manuscript; I.B., M.G., G.H., N.M., and A.C. conducted experiments and data analyses; K.A.R. and S.S. performed bioinformatics analyses; I.B. and N.M. generated transgenic lines; C.R.A contributed to interpretation of the results, provided important intellectual content and co-wrote the manuscript; K.Y. directed the study, secured funding, designed experiments, analyzed data and co-wrote the paper with inputs from all authors.

The authors declare no competing interests.

## Materials and Methods

### Zebrafish husbandry, transgenic and mutant lines

Zebrafish were raised by standard methods and handled according to the guidelines of the Weizmann Institute Animal Care and Use Committee^73^. For all imaging, *in situ* hybridization, and immunofluorescence experiments, embryos were treated with 0.003% phenylthiourea (PTU, Sigma-Aldrich) from 8 hpf to inhibit pigment formation. Zebrafish lines used in this study were: *Tg(fli1:EGFP)^yl^* ^74^*; Tg(gata1a:DsRed)^sd2^* ^74^*; Tg(kdrl:EGFP)^s843^* ^74^; *Tg(kdrl:mCherry)^y206^* ^75^; *Tg(gfap:Tomato)^nns17^* ^76^; *Tg(gfap:GAL4FF)^wa32^* ^77^; *Tg(olig2:EGFP)^vu12^* ^78^; *Tg(olig2:dsRed)^vu19^* ^78^; *Tg(isl1a:EGFP)^rw0^* ^78^; *Tg(slc1a3b:MYRGFP-2A-H2AmCherry)*^79^; *Tg(EPV.Tp1-Mmu.Hbb:EGFP)^ia12^* ^44^; *Tg(kdrl:TagBFP)^mu293^* ^80^; *Tg(glut1b:mCherry)^sj1^* ^81^; *Tg(mbp:EGFP)^ck1^* ^32^; *Tg(hsp70I:N3ECD-EGFP)^co15^* ^82^; *Tg(hsp70I:N3ICD-EGFP)^co17^* ^82^; *Tg(hsp70l:MYC-notch1a,cryaa:Cerulean)^fb12^* ^57^; *notch3^fh332/fh332^* ^82^, *Tg(hsp70l:lamp1b-RFP)^pd1064^* ^83^, *Tg(h2ax:EGFP-rab7a)^mw7^* ^84^.

The *TgBAC(scar2ba:KalTA4)* line was generated using the DKEY-177l10 BAC (BioScience, HUKGB735J035Q) containing the full *scarb2a* sequence. The KalTA4 fragment was cloned downstream to the ATG following the removal of the *scarb2*a CDS. The resulting BAC*(scar2ba:KalTA4)* construct was injected at the 1-cell stage into AB zebrafish and integrated through transposon-mediated BAC transgenesis to generate a stable reporter line that was then crossed with *scarb2a* mutant fish^85^.

*Tg(UAS-scarb2a-P2A-RFP)* and *Tg(UAS-scarb2a;cmcl2:EGFP)* lines were generated by amplifying the coding sequences of *scarb2a* with the following primers: F 5’-ATGACTAGAAGATCTTGTACTATTTACGCC-3’; R 5’-TCAACACTTTTGTGCCTCCACT-3’.

The resulting fragment was then cloned in pDONR221 middle donor plasmid and combined with p5E-UAS, p3E-P2A-mKate2, and pDestTol2pA2 using the Gateway system as previously described^86^.

The *scarb2a* mutant was generated using CRISPR/CAS technology. The CRISPR guide was designed with CHOPCHOP (https://chopchop.cbu.uib.no/), and potential off-target sequences were assessed using the MIT CRISPR Design site. The guide sequence GGCTGAGAACAGCAGAGTAT was cloned into the BsmBI sites of the pT7-gRNA plasmid (Addgene plasmid 46759). Cas9 mRNA was generated from pCS2-nCas9n^87^ using the mMACHINE T7 ULTRA kit (Ambion, AM1345). Cas9 mRNA (250 ng/μL) and guide RNA (100 ng/μL) were co-injected into 1-cell-stage embryos. For genotyping, genomic DNA was extracted and amplified with 5′-TGTCTGTGTTTGGTTACAGGAG-3’ and 5′-TTCCCCGCCAAGAACTCA-3’ (138 bp) primers and analyzed on a 2% agarose gel.

### Angiography

Tetramethylrhodamine Dextran (molecular mass, 2,000,000 Da; Thermo Fisher Scientific, D7139) was injected intravascularly on anesthetized *larvae* as previously described^88^. Imaging was initiated within 5 min of the injection.

### *In situ* hybridization and immunostaining

*In situ* hybridization was performed as described^89^. The following primers were used to generate the *glut1b* riboprobes: 5′-ATTGGCATCCTCATGGCACA-3’; 5’-ATGAAAACGTATGGGCCGGT-3’

For immunostaining, 60 hpf embryos were fixed in 4% PFA overnight, permeabilized in acetone for 25 min at RT, blocked in PBS containing 2% BSA, 5% goat serum, and 0.1% Tween20 for 3 hrs. at RT, and incubated overnight at 4°C with rabbit anti-Sox2 (1:100; Abcam ab97959) or mouse anti-HuC (1:400; Thermo Fisher Scientific A-21271) antibodies. After extensive washing with PBST, embryos were incubated overnight at 4°C with secondary antibodies conjugated with Alexa Fluor 633 (1:500; Goat anti-Rabbit Invitrogen A21070, Goat anti-Mouse Invitrogen A21053).

### Lysotracker staining

Lysotracker™ Deep Red (Thermo Fisher L12492) staining was previously described^90^. Briefly, 25-30 embryos were transferred into 6-well plates containing LysoTracker™ Deep Red dissolved to a final concentration of 10 μM in PTU water and incubated at 28.5°C for 45 min in a dark environment. Embryos were then rinsed 3 times with ∼1 ml of fresh fish water and imaged immediately using a 633 nm laser for far red staining. The number of LysoTracker-positive punctae per unit area was counted.

### Heat shock induction

Heat shock was performed at 37 °C for 45 minutes. For *Tg(hsp70I:N3ECD-EGFP)^co15^*, *Tg(hsp70I:N3ICD-EGFP)^co17^* and *Tg(hsp70l:MYC-notch1a,cryaa:Cerulean)^fb12^* experiments, the heat shock was repeated every 6-7 hours on embryos from 28 hpf until 60 hpf.

### Drug Treatments

To inhibit γ-secretase, 28 hpf manually dechorionated embryos were incubated at 28.5 °C in the dark into a 2 ml petri dish containing 0.4% DMSO in embryo water with LY-411575 (Sigma cat. SML0506) at a final concentration of 50 μM. To specifically block V-ATPase proton pump activity, we incubated 28 hpf zebrafish embryos with Bafylomicin A1 inhibitor (Sigma cat. B1793) that was added directly to the fish medium at a final concentration of 100 nM. 0.4% DMSO was used as control. In all treatments, the embryos were left in the incubation medium at 28.5 °C until 60 hpf.

### Cell sorting and Bulk MARS-Seq

For each experimental condition, a pool of 50 heads (3 biological replicates) was dissected from euthanized 48 hpf *scarb2a* mutants and wt siblings and dissociated into a single-cell suspension. After a brief enzymatic treatment with a cocktail of Liberase Blendzyme 3 (Roche), trypsin B (BI), and DNAseI (Roche), the cell suspension was strained through a 70µm filter and stained with SYTOXTM blue (ThermoFisher) for live/dead discrimination. 5000 live *scarb2a*+ single cells were sorted into 40 µl of lysis/binding buffer solution (ThermoFisher) containing RNase inhibitor RNasinTM (Promega). FACS analysis and sorting were performed on a (BD FACS Aria III) using a 70µm nozzle. RNA was captured using Dynabeads™ mRNA DIRECT™ Purification Kit (ThermoFisher) prior to library preparation. A bulk adaptation of the MARS-seq protocol^91^ was used to generate RNA libraries for the expression profile of *scarb2*+ mutant and wt cells. The RNA was further fragmented and transformed into a sequencing-ready library by tagging the samples with Illumina sequences during ligation, RT, and PCR. The final library concentration was measured by Qubit, TapeStation, and qPCR for zebrafish actin as previously described^91^. Sequencing was performed on a Nextseq500/550 High Output Kit v2.5 75 cycles (Illumina; paired-end sequencing), and each sample was sequenced for 6M reads. The Initial quality control report was done using the User-friendly Transcriptomic Analysis Pipeline^92^. Upregulated and downregulated transcription factors were identified in the sequenced dataset using gene ontology from Uniprot.org.

### Total RNA isolation and quantitative reverse transcription PCR

A pool of 10–20 heads/sample was dissected from 48 hpf wt and mutant embryos. The samples were homogenized in TRIzol (Invitrogen, Thermo Fisher Scientific, 15596026) and processed for RNA extraction following standard procedures^88^. A total of 1 μg of RNA per reaction was reverse-transcribed using a High-Capacity cDNA Reverse Transcription Kit (Applied Biosystems, Thermo Fisher Scientific, 4368814). The following primers were used:

*notch1a* 5′-GTGGCACAAGGGCAAAGATG -3′, 5′-CAGGGGTTCGGGAATTGACA-3′;

*notch3* 5′-GCATTGACCGACCTAATGGA -3′, 5′-TGCTCTCACACAGTCTTCCTTC-3′;

*her4.2* 5’-GGCTCAATCAGCAGCAGAGA-3’; 5’-GCACTGCTTTTCTGAGAGCG-3’;

her6 5’-AATGACCGCTGCCCTAAACA-3’; 5’-TCACATGTGGACAGGAACCG-3’.

Expression levels were standardized to the primer set specific for β-actin 5′-TGACAGGATGCAGAAGGAGA-3′ and 5′-GCCTCCGATCCAGACAGAGT-3′.

### Microscopy and imaging

Confocal imaging was performed using a Zeiss LSM780 upright confocal microscope or LSM880 upright confocal microscope equipped with an Airyscan module 32-channel GaAsP-PMT area detector, water-immersed ×20/1.0 NA or ×10/0.5 NA objective lens. Embryos were mounted using 1.5% (w/v) low-melting agarose.

### Image processing

Confocal images were processed offline using the Fiji^93^ version of ImageJ (NIH) or Imaris v.9.3 (Bitplane). The images shown in this study are single-views, 2D-reconstructions of collected *z*-series stacks. The co-localization channel was created using the Imaris Colocalization Module. Co-localization thresholds were set manually. Transversal sections were prepared using Imaris 3D software from z-series stacks. Detection, colocalization, and quantification of Lysotracker, Lamp1, and Rab7 positive puncta were performed using ComDet v.0.5.5 plugin for ImageJ (https://github.com/UU-cellbiology/ComDet).

### In silico analyses

For Extended Data Fig. 1a, the amino acid sequences for zebrafish (Q8JGR8, 531aa), mouse (O35114, 478 aa), and human (Q14108, 478 aa) were obtained from Uniprot.org. Percentage identity were performed using Uniprot.org.

For Extended Data Fig. 2a, previously published single-cell RNA sequencing data from developing mouse brain and spinal cord were used^94^. Bar graphs were made using R (version 4.3.1) with the ggplot2 package^95^.

For Extended Data Fig. 3a, single-cell transcriptomic data from 310 cells isolated from *Tg(olig1:memEYFP)* zebrafish at 5 dpf were obtained from^96^. Plots were obtained from a searchable database made publicly available at https://castelobranco.shinyapps.io/zebrafish_OPCs/ ^96^

### Statistical analyses

Statistical significance between two samples was calculated using unpaired two-tailed Student’s *t*-tests assuming unequal variance from at least three independent experiments unless stated otherwise. Ordinary one-way ANOVA with Tukey’s multiple comparisons was used when comparing more than two groups to test mean differences. In all cases, normality was assumed, and variance was comparable between groups. Sample size was selected empirically according to previous experience in assessing experimental variability. The investigators were blind to allocation during experiments and outcome assessment. Numerical data are the mean ± s.e.m. unless stated otherwise. Statistical calculations and the graphs for the numerical data were performed using Prism 8 software (GraphPad Software).

