## Extended Data for "Endolysosomal dysfunction in radial glia progenitor cells leads to defective cerebral angiogenesis and compromised Blood-Brain Barrier integrity"

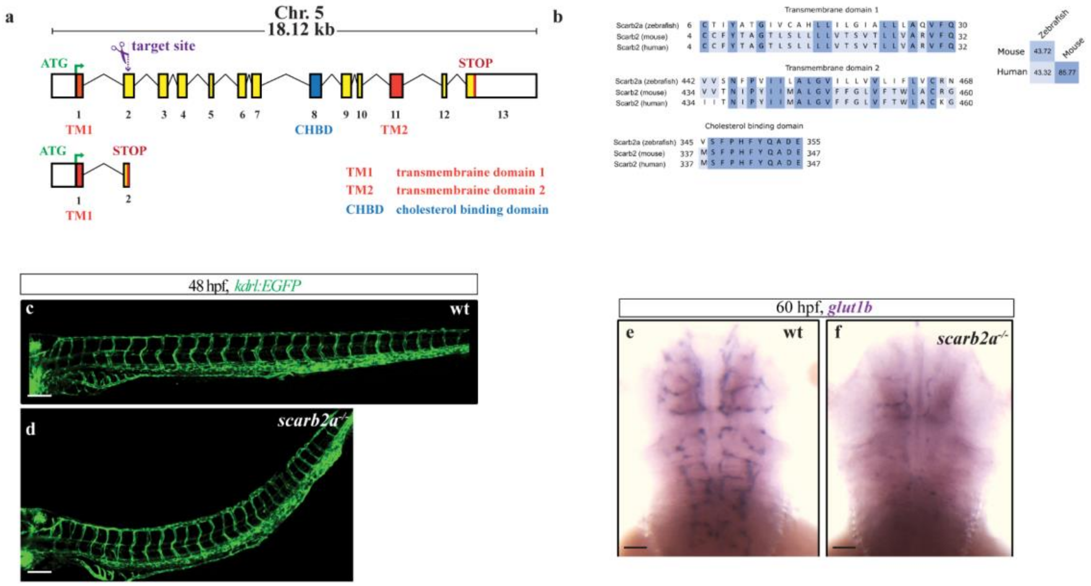


**Extended Data Fig. 1: *scarb2a* is homologous to mammal SCARB2, and its mutation exclusively affects the brain vasculature. a**, Schematic illustration showing *scarb2a* wt coding sequence with target sites of guide RNAs (gRNA, purple) and resulting *scarb2a* mutant sequence. **b**, Conserved transmembrane domains and cholesterol binding domains in zebrafish (Scarb2a), mouse (SCARB2), and human (SCARB2). Next to the comparison is the percentage identity matrix for SCARB2 aminoacid sequences between the species. **c-d**, Lateral views of *Tg(kdrl:EGFP)* in wt (c) and *scarb2a*$^{-/-}$ (d) embryos at 48 hpf showing no defects in trunk vasculature of mutant embryos. **e-f**, Whole mount ISH against *glut1b* at 60 hpf (purple, dorsal view). TM1, transmembrane domain 1; TM2, transmembrane domain 2; CHBD, cholesterol binding domain. Scale bars, d-e=200 µm; f-g=50 µm.

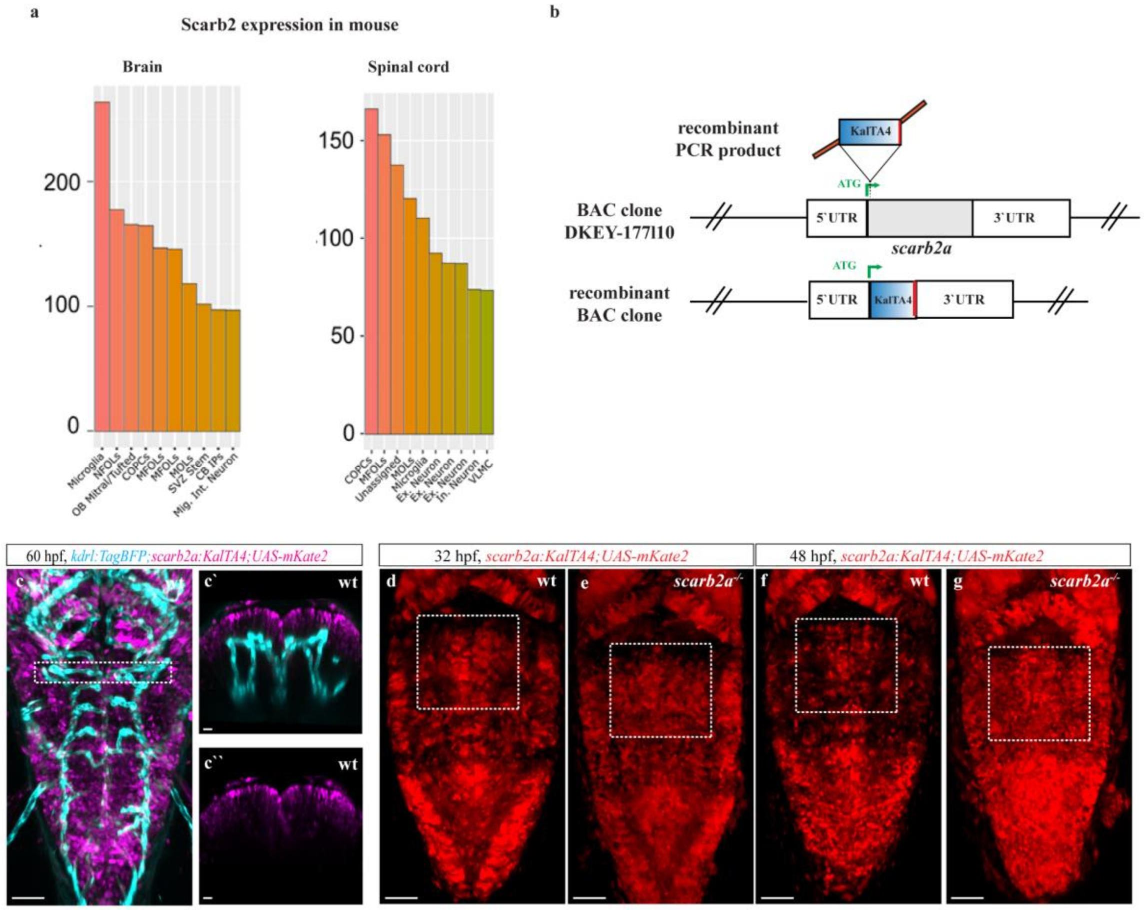


**Extended Data Fig. 2: *scarb2a* is expressed in RGC progenitors. a**, Top 10 cell types with higher expression of Scarb2 in the developing mouse brain and spinal cord (Single-cell transcriptomics dataset re-analyzed from[19]). **b**, Illustration of the construct used to generate *TgBAC(scarb2a:KalTA4)* reporter. **c**, Dorsal view of *Tg(kdrl:TagBFP;scarb2a:KalTA4;UAS-mkate2)* (*scarb2a* in magenta) in wt embryos at 60 hpf (dashed square in c marks the region shown in **c'** and **c''**) showing no colocalization of *scarb2a* and *kdrl* (blue). **d-g**, Selected confocal images from *Tg(scarb2a:KalTA4;UAS-mkate2)* (Extended Data Video 2) at 32 (d,e) and 48 hpf (f,g). Scale bars, c',c''=30 µm; c,f-g= 60 µm.

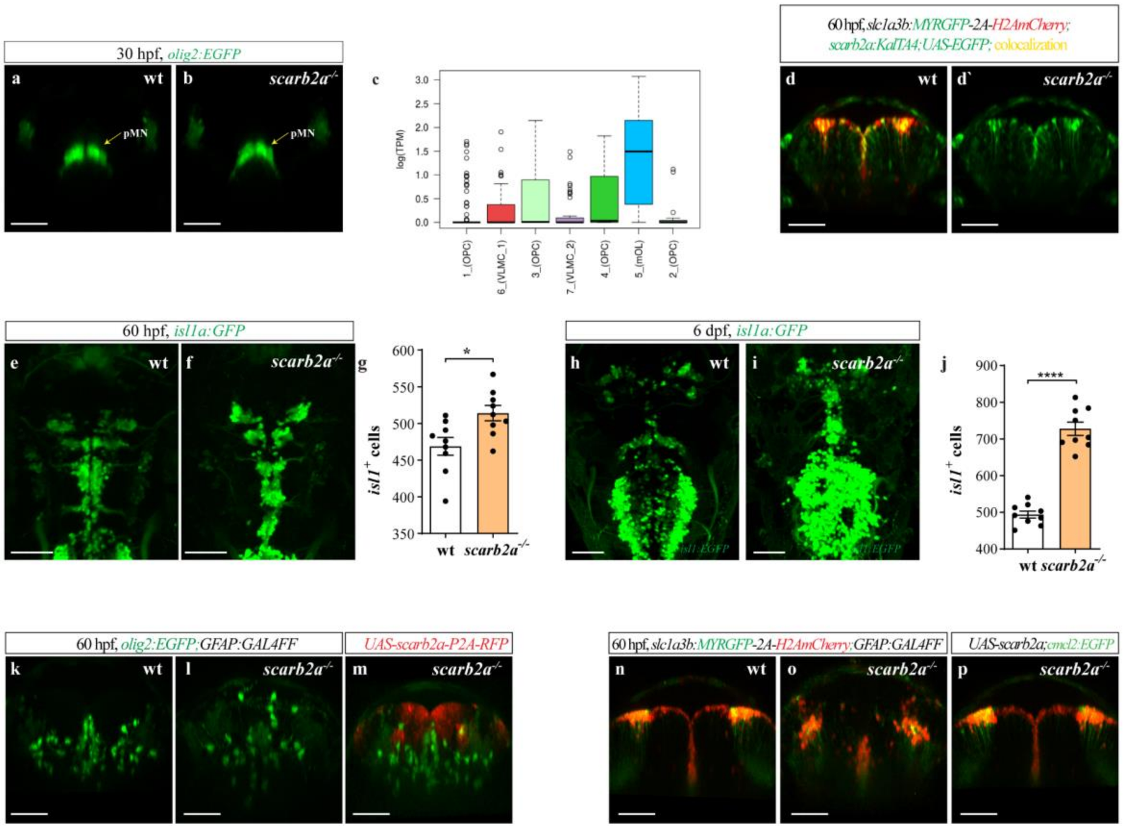


**Extended Data Fig. 3: *Scarb2a* depletion leads to abnormal gliogenesis a-b**, Transverse optical section at the level of the pMN in *Tg(olig2:EGFP)* wt (a) and *scarb2a* mutant (b). **c**, Analysis of single-cell transcriptomic data obtained from[33] (Oligodendroctytes precursors (OPC) = 1_OPC, 2_OPC, 3_OPC, 4_OPC; mature oligodendrocytes (mOLs) = 5_mOL, vascular and leptomeningeal cells (VLMCs) = 6_VLMC_1, 7_VLMC_2. **d-d`**, Transverse optical section of *Tg(slc1a3b:MYRGFP-2A-H2AmCherry scarb2a:KalTA4;UAS-EGFP)* double transgenic reporter. **e-j,** Dorsal views of *Tg(isl1:EGFP)* reporter showing increased numbers of *isl1*$^{+}$ neurons in mutant hindbrains at 60 hpf (e-g, n =9, two-tailed Student`s *t*-test) and 6 dpf (h-j, n =9, two-tailed Student`s *t*-test add details). **k-m**, Transverse optical sections of *Tg(olig2:EGFP;GFAP:Gal4FF)* in wt (k), *scarb2a*$^{-/-}$ (l), and *scarb2a*$^{-/-}$ mutant following injection of *UAS-scarb2awt-P2A-RFP* construct (m, red). **n-p,** Transverse optical sections of *Tg(slc1a3b:MYRGFP-2A-H2AmCherry; GFAP:Gal4FF)* in wt (n), *scarb2a*$^{-/-}$ (o) and *scarb2a*$^{-/-}$ injected with *UAS-scarb2a,cmcl2:EGFP* construct (p).. Scale bars, a-b, d-d`, k-m, n-p=30 µm; e-f, h-i=50 µm. Error bars are mean ± s.e.m.

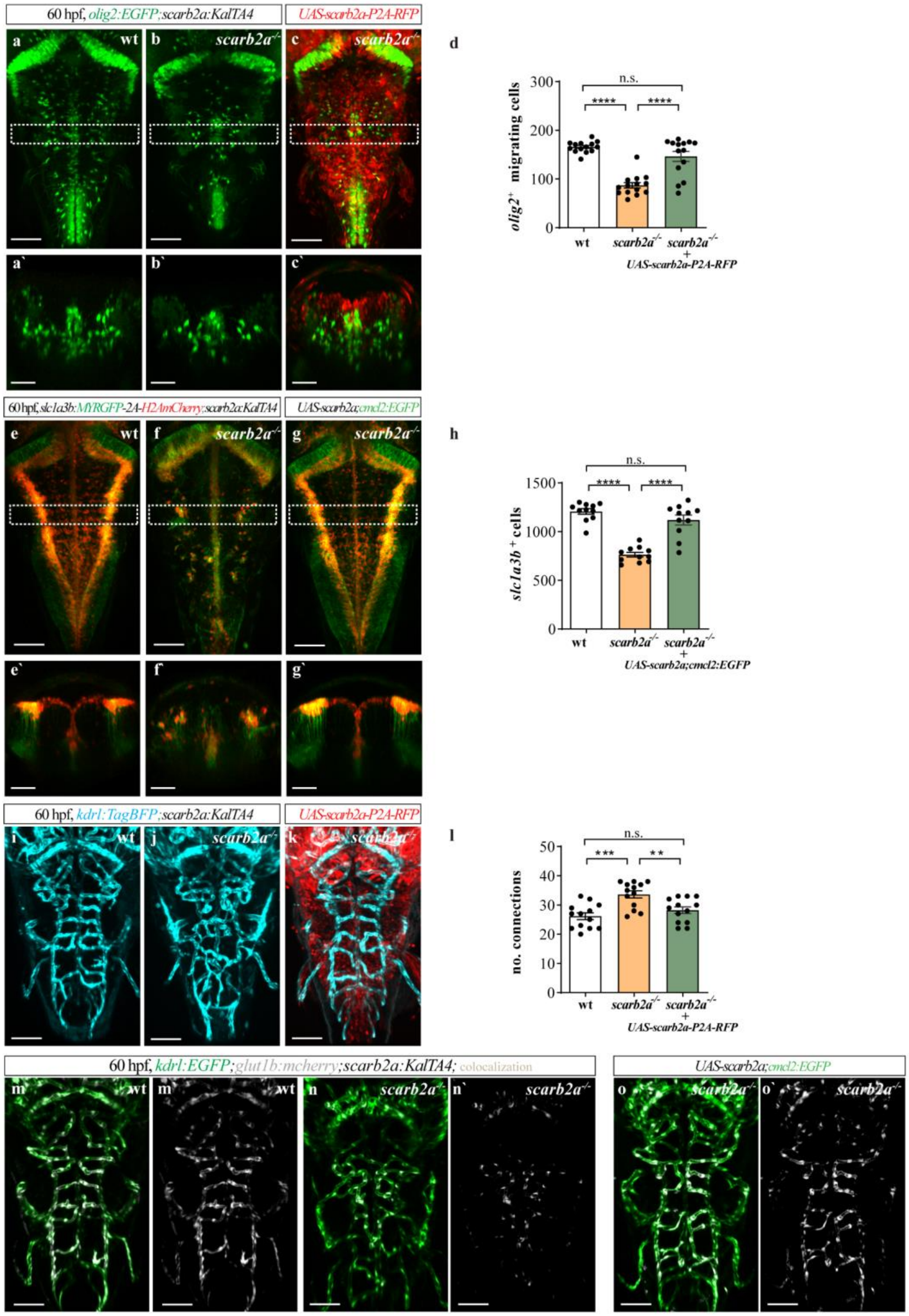

60 hpf, olig2:EGFP;scarb2a:KalTA4
UAS-scarb2a-P2A-RFP
a wt
b scarb2a-/-
c scarb2a-/-
d
olig2+ migrating cells
n.s.
****
****
wt
scarb2a-/-
scarb2a-/- + UAS-scarb2a-P2A-RFP
a`
b`
c`
60 hpf, slc1a3b:MYRGFP-2A-H2AmCherry;scarb2a:KalTA4
UAS-scarb2a;cmcl2:EGFP
e wt
f scarb2a-/-
g scarb2a-/-
h
slc1a3b+ cells
scarb2a-/- + UAS-scarb2a;cmcl2:EGFP
e`
f`
g`
60 hpf, kdrl:TagBFP;scarb2a:KalTA4
i wt
j scarb2a-/-
k scarb2a-/-
l
no. connections
***
**
60 hpf, kdrl:EGFP;glut1b:mcherry;scarb2a:KalTA4; colocalization
m wt
m` wt
n scarb2a-/-
n` scarb2a-/-
o scarb2a-/-
o` scarb2a-/-

**Extended Data Fig. 4: Restoring Scarb2a expression in *scarb2a*+ cells results in complete recovery of neuronal, glial and vascular phenotype a-c`**, Dorsal views (a-c) and transverse optical sections (a`-c`) of *Tg(olig2:EGFP;scarb2a:KalTA4;)* at 60 hpf. Injection of *UAS-scarb2a-P2A-RFP* led to a complete rescue of OPCs in *scarb2a*$^{-/-}$ embryos (c-c`). **d**, Quantification of migrating *olig2*$^{+}$ cells in a-c (n=14; One-way ANOVA, multiple comparisons with Tukey posthoc test). **e-g'**, Injection of *UAS-scarb2awt,cmcl2:EGFP* construct into *Tg(slc1a3b:MYRGFP-2A-H2AmCherry;scarb2a:KalTA4)*;*scarb2a*$^{-/-}$ results in full recovery of the astrocyte population (g-g'). **h**, Quantification of *slc1a3b*$^{+}$ red nuclei in e-g (n= 11; One-way ANOVA, multiple comparisons with Tukey post-hoc test). **i-k**, Dorsal views of *Tg(kdrl:TagBFP;scarb2a:KalTA4)* hindbrains in wt (i), *scarb2a*$^{-/-}$ (j) and *scarb2a*$^{-/-}$ following *UAS-scarb2a-P2A-RFP* injection (k) embryos. **l**, Quantification of CtA interconnections in i-k (n= 13; One-way ANOVA, multiple comparisons with Tukey posthoc test). **m-o`**, Dorsal views of *Tg(kdrl:EGFP;glut1b:mCherry;GFAP:Gal4FF)* depicting complete recovery of *glut1b* expression (white) following injection of *UAS-scarb2a,cmcl2:EGFP* in mutant embryos (o-o`). Scale bars: a-c, e-g, i-k, m-o'=60 μm; a'-c', e'-g'= 30 μm. Error bars are mean ± s.e.m.

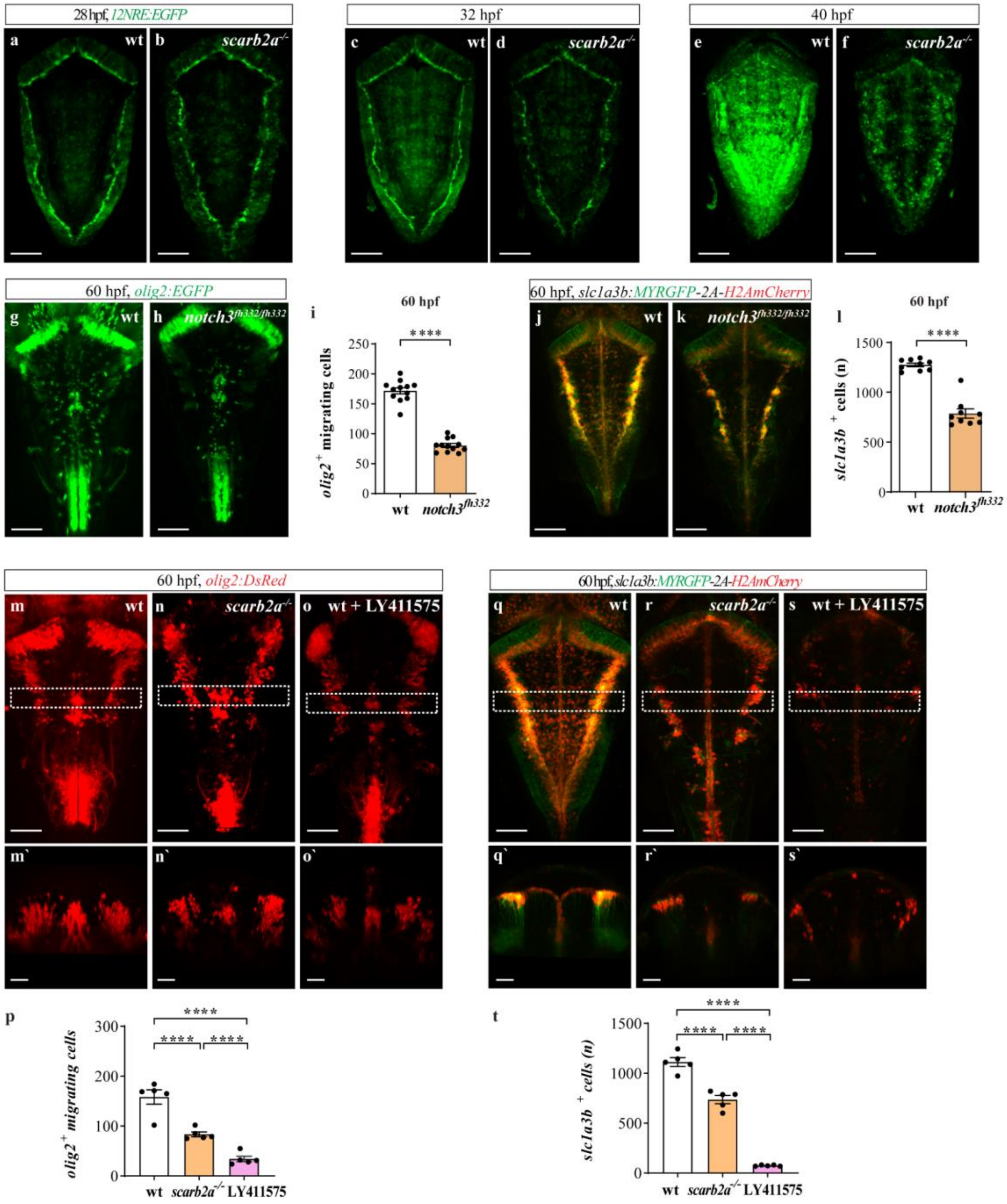
28 hpf, 12NRE:EGFP
32 hpf
40 hpf
a wt
b scarb2a-/-
c wt
d scarb2a-/-
e wt
f scarb2a-/-
60 hpf, olig2:EGFP
g wt
h notch3fh332/fh332
i
60 hpf
****
olig2+ migrating cells
250
200
150
100
50
0
wt
notch3fh332
60 hpf, slc1a3b:MYRGFP-2A-H2AmCherry
j wt
k notch3fh332/fh332
l
60 hpf
****
slc1a3b + cells (n)
1500
1000
500
0
wt
notch3fh332
60 hpf, olig2:DsRed
m wt
n scarb2a-/-
o wt + LY411575
60 hpf, slc1a3b:MYRGFP-2A-H2AmCherry
q wt
r scarb2a-/-
s wt + LY411575
m`
n`
o`
q`
r`
s`
p
****
****
****
olig2+ migrating cells
300
200
100
0
wt
scarb2a-/-
LY411575
t
****
****
****
slc1a3b + cells (n)
1500
1000
500
0
wt
scarb2a-/-
LY411575

**Extended Data Fig. 5: Notch signaling downregulation in *scarb2a*$^{-/-}$ lead to similar phenotype of *notch3*$^{fh332}$ mutant**

**a-f**, Dorsal views of *Tg(12NRE:EGFP)* reporter, at 28 (a-b), 32 (c,d) and 40 (e,f) hpf depicting significant reduction of Notch signaling activation in mutant hindbrains starting at 32 hpf (d,f). **g-i'**, Confocal images of *Tg(olig2:EGFP);notch3*$^{fh332/fh332}$ hindbrains showing significant decrease of OPC numbers (h) as compared to wt siblings (g) (dashed square in g-h, marks the region shown in g'-h'), quantified in (i) (n=12, two-tailed Student`s *t*-test, *P*<0.0001). **j-l**, Confocal images of *Tg(slc1a3b:MYRGFP-2A-H2AmCherry);notch3*$^{fh332/fh332}$ revealed an absence of *slc1a3b*$^+$ cells in the vz of *notch3* mutant (k) (dashed square in j-k, marks the region shown in j'-k'); quantified in (l) (n=9, two-tailed Student`s *t*-test, *P*<0.0001). **m-o`**, Dorsal views of *Tg(olig2:DsRed)* (m-o') embryos showing defects in migrating OPCs following LY411575 treatment, quantified in (p) (n=5, One-way ANOVA, multiple comparisons with Tukey post-hoc test). **q-s**', Dorsal views of *Tg(slc1a3b:MYRGFP-2A-H2AmCherry)* embryos showing absent *slc1a3b*$^+$ cells following LY411575 treatment, quantified in (t), (n=5, One-way ANOVA, multiple comparisons with Tukey posthoc test). Scale bars: a-f= 50 μm; g-h, j-k, m-o, q-s=60 μm; m'-o', q'-s'=30 μm. Error bars are mean ± s.e.m.

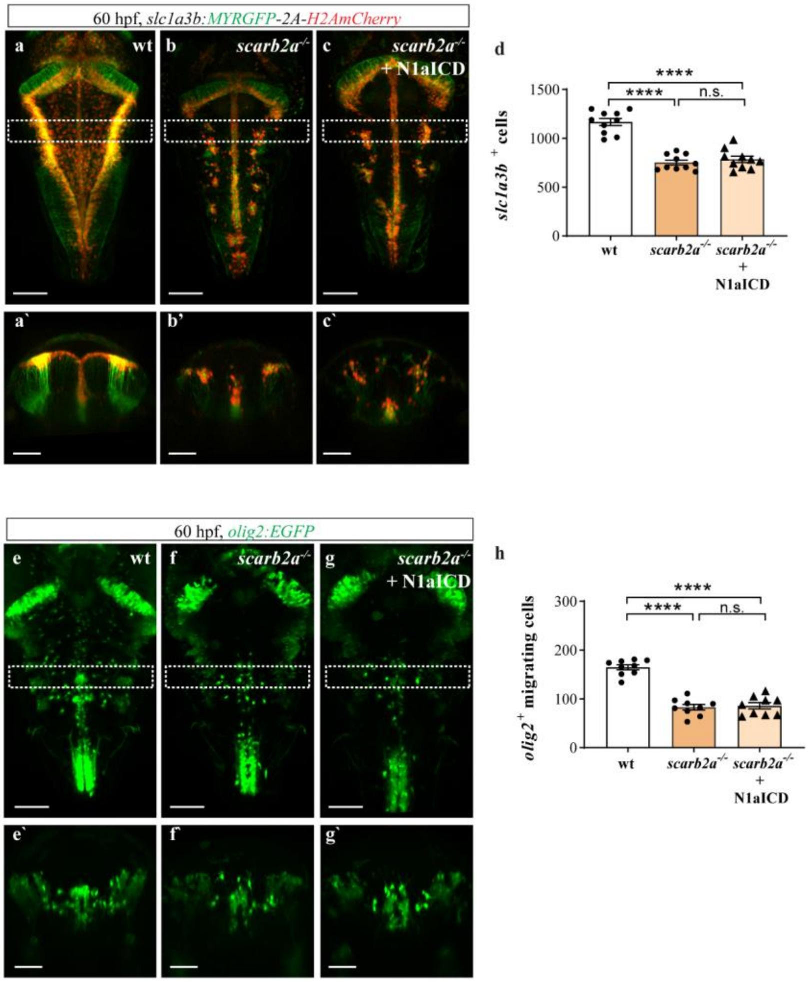


**Extended Data Fig. 6: Overexpression of Notch1a intracellular domain (N1aICD) does not rescue *scarb2a* gliogenic phenotype a-c`**, Confocal images of heat-shock induced overexpression of *notch1* intra-cellular domain (N1aECD) in *Tg(slc1a3b:MYRGFP-2A-H2AmCherry;hsp70l:MYC-notch1a-cryaa:Cerulean)* mutant embryos (dashed square in a-c, marks the region shown a'-c'). **d,** Quantification of *slc1a3b*$^+$ astrocytes in the hindbrain of a-c (n=10, One-way ANOVA, multiple comparisons with Tukey posthoc test). **e-g**`, Dorsal views of *Tg(olig2:EGFP;hsp70l:MYC-notch1a,cryaa:Cerulean)*;*scarb2a* embryos with or without heat-shock induced overexpression of N1ICD. **h**, Quantification of hindbrain *olig2*$^+$ migrating OPCs (n=9, One-way ANOVA, multiple comparisons with Tukey post-hoc test). Scale bars: dorsal views a-c, e-g= 60 μm; a'-c', e'-g'=30 μm. Error bars are mean ± s.e.m.
